# Genomes of keystone *Mortierella* species lead to better *in silico* prediction of soil mycobiome functions from Taiwan’s offshore islands

**DOI:** 10.1101/2021.12.20.473467

**Authors:** Yu-fei Lin, Wei-An Liu, Yu-Ching Liu, Hsin-Han Lee, Yen-Ju Lin, Ed-Haun Chang, Meiyeh J Lu, Chih-Yu Chiu, Isheng Jason Tsai

## Abstract

The ability to correlate the functional relationship between microbial communities and their environment is critical to understanding microbial ecology. There is emerging knowledge on island biogeography of microbes but how island characteristics influence functions of microbial community remain elusive. Here, we explored soil mycobiomes from nine islands adjacent to Taiwan using ITS2 amplicon sequencing. Geographical distances and island size were positively correlated to dissimilarity in mycobiomes, and we identified 56 zero-radius operational taxonomic units (zOTUs) that were ubiquitously present across all islands, and as few as five *Mortierella* zOTUs dominate more than half of mycobiomes. Correlation network analyses revealed that seven of the 45 hub species were part of the ubiquitous zOTUs belonging to *Mortierella, Trichoderma, Aspergillus, Clonostachys* and *Staphylotrichum*. We sequenced and annotated the genomes of seven *Mortierella* isolates, and comparative predictions of KEGG orthologues using PICRUSt2 database updated with new genomes increased sequence reads coverage by 62.9% at the genus level. In addition, genes associated with carbohydrate and lipid metabolisms were differentially abundant between islands which remained undetected in the original database. Predicted functional pathways were similar across islands despite their geographical separation, difference in differentially abundant genes and composition. Our approach demonstrated the incorporation of the key taxa genomic data can improve functional gene prediction results and can be readily applied to investigate other niches of interests.

## 1. Introduction

Fungi are one of the most diverse groups of organisms in the biological kingdom— with an estimated 6.2 million species [1]—and play an essential in ecosystems with their ability to decompose organic matters encompassing all ecological niches, from soil [2] to water [3]. The Earth Microbiome Project was the catalyst for an extensive profiling of microbes in soil [4–7]. Protocols for metabarcoding characterisation of eukaryotic species quickly followed [8,9], for instance, characterising the fungal community of a given environment is defined as the mycobiome [10]. Climate and vegetation were determined to be the main factors driving mycobiome community diversity and structure [11,12]. Profiling of soil biomes around the world showed that earth’s soil biome is dominated by as few as 83 fungal phylotypes. The predominant soil fungal phylum is Ascomycota, accounting for ∼18% of sequence abundance [13].

Although recent efforts to characterise the distribution and abundance of fungal species in different niches have revealed insights around host-fungal relationships [14–16], little is known about the functional relationship between fungal communities and their respective niches. Metatranscriptomics is a direct approach to elucidating the relationship between the fungal community and its surroundings. For example, the ectomycorrhizae community metatranscriptome and its metabolic pathways have been shown to respond to perturbations caused by different fertilisation strageties in Norwegian spruce trees [17]. Unfortunately, a major technical hurdle in metatranscriptomic approach is the strong bias in the host-fungi biomass ratio [18]. Challenges include i) most sequencing reads from samples originated from the host, making only a few fungal reads available for subsequent analyses, and ii) filtering of reads from the host might not be readily applicable due to the limited availability of the host genome. In addition, not enough fungal genomes are available from various environments, rendering challenges in classification and subsequent analyses of fungal sequences.

To overcome these limitations, an alternative and cost-effective approach to delineating microbial functional relationships is *in silico* inference using tools—e.g., Tax4Fun [19], FAPROTAX [20] and PICRUSt2 [21]—developed to infer functions of the microbiome from relative abundances of phylogenetically classified amplicons. These tools have been used extensively, especially in many gut bacterial microbiome studies. Functional changes associated with perturbations in the gut microbiome from antibiotics [22] or disease states [23] have been predicted. Benchmark studies show that predictions made from amplicon sequencing are comparable to those from shotgun sequencing of four independent sets of microbiome data, with similar accuracy [21], which is the current gold standard for inferring functional gene family and pathway [24–26]. Such tools also exist for mycobiomes, such as FUNGIpath, which reconstructs fungal metabolic pathways by predicting putative pathways from protein sequence orthologies [27]. Annotated fungal genomes allow for further categorisation into functional guilds based on their trophic state using FUNGuild [28], reflecting their putative ecological roles. *In silico* prediction of functions remained limited in fungi owing to limited genomic data. For example, PICRUSt2 utilises 41,926 bacterial and archaeal genomes, but only 190 fungal genomes [21], and is biased towards model organisms [29]. One major fungal taxon that lacks genome information is the genus *Mortierella*, which is ubiquitous in soil samples [11,30]; the GlobalFungi database (https://globalfungi.com, accessed September 2021) revealed that *Mortierella* can be found in all the deposited soil sample records (n=18,759). However, there are only 44 publicly available *Mortierella* genomes (16 and 28 deposited in the JGI MycoCosm and NCBI databases, respectively). This genus has recently received attention as it was shown to be an important ectomycorrhiza for facilitating plant development. There is currently an urgent need to incorporate well-annotated genomes to yield better inferences on mycobiomes’ functions across various niches.

This study aims to characterise the functional roles of mycobiomes in forests of nine offshore islands proximal to Taiwan. We examined fundamental features of island biogeography—e.g., the species-area distribution relationship, species vicariance and the distance-decay relationship [31,32]. We identified the key fungal taxa to be the *Mortierella* genus, then isolated and sequenced the taxa to obtain high-quality and annotated genomes. We assessed whether functional metabolic pathway predictions of the mycobiome could be improved by incorporating novel genomes into existing functional pathway prediction workflows from amplicon studies and compared them with the current pipeline.

## 2. Materials & methods

### 2.1 Study sites and sample collection

The study was conducted on several remote islands **(Fig.1)**. The archipelagos of Matsu islands (MT), are located 10–50km offshore of mainland China and face the Taiwan Strait, including Beigan (MT-BG), Nangan (MT-NG), Dongju (MT-DJ), Hsiju (MI-SJ), and Dongyin (MT-DY) Islet. Two tropical volcanic islands, Orchid Island (OI), and Green Island (GI) are located about 60 and 30 km, respectively, from the southeastern part of Taiwan and face the Pacific Ocean. The coral reef-originated Dongsha (DS) Islet and Taiping (TP) Islet are located south-west of Taiwan in the South China Sea.

**Fig. 1.**
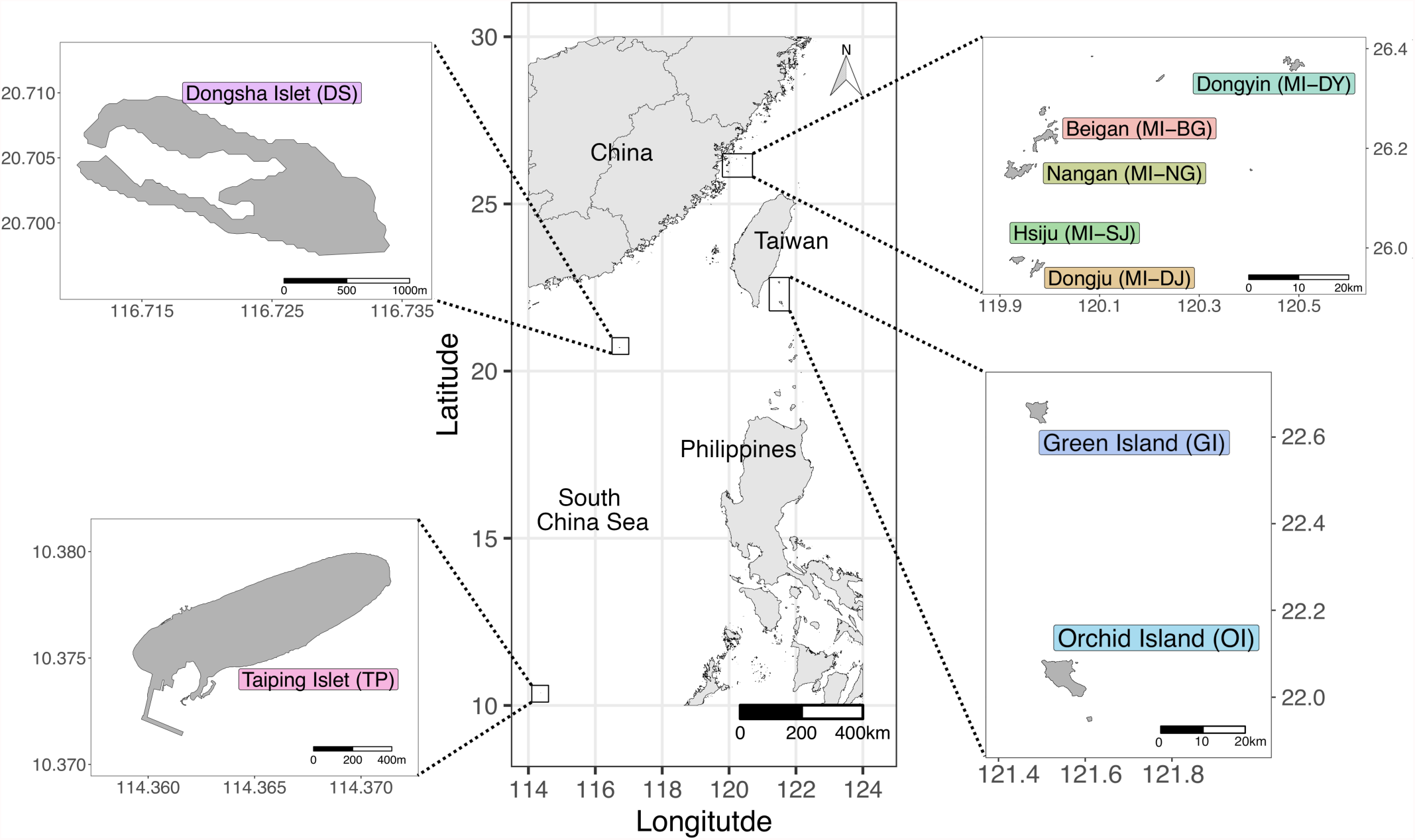
Map of the sampling sites from Taiwan’s offshore islands.

The soils on the five MT islands comes from granite parent material, and classified as haplustults. The soils on OI and GI come from andesite parent material, and were classified as paleudults. Detail of soil and forest types was described in [33,34]. The soil on DS and TP are classified as entisol, and their native vegetation are predominantly covered with screw pine (*Pandanus tectorius*) and natural tropical forest, respectively.

From 2016–2017, surface soil samples (0–10 cm deep) from the MT, OI and GI were collected as previously described by Lin et al. [33]. Soil from the DS and TP were sampled in 2018–2019 (**Fig. 1**). Leaf litter was avoided. Approximately ten to twelve 10 cm deep soil cores were collected using a core borer. Soil samples were stored on ice before transporting back to the laboratory, where they were sieved through 2mm steel mesh and homogenised by manual mixing and stored at −20°C until use. Edaphic data were collected as described previously [33,34], with the following metadata collected: microbial biomass carbon (MBC), microbial biomass nitrogen (MBN), microbial biomass phosphate (MBP), pH, cellulase, xylanase, β-glucosaminidase, phosphomonoesterase, urease, proteinase, total phospholipid-derived fatty acid (PLFA), fungal PLFA, bacterial PLFA, arbuscular mycorrhizal fungal PLFA (AMF-PLFA), and the main vegetation around the sample site. Mean monthly temperature (MMT) and mean monthly precipitation (MMP) were taken from the records of the nearest weather station (https://opendata.cwb.gov.tw/index).

### 2.2 Total genomic DNA extraction from soil samples

Genomic DNA of homogenised soil was extracted using PowerSoil DNA extraction kits (Qiagen, Hilden, Germany). Approximately 0.25 g of soil was extracted from each sample with minor modifications to the manufacturer’s protocol. Sample homogenisation and lysis were performed using PowerLyzer 24 (Qiagen, Hilden, Germany) set at 2,000 rpm for 2 × 5 min. After solutions C2 and C3 were added, the lysate were kept at 4° for 1 hr.

### 2.3 Recovery of *Mortierella* isolates from soil

To isolate *Mortierella* species, one gram of soil sample was serially diluted ten-fold in 1.0 mL up to 10^−5^ with 1 X PBS. For each dilution, 200 μL of suspension was plated onto potato dextrose agar (PDA) supplemented with 50 μg/mL chloramphenicol and incubated at 28°C and monitored daily for fungal growth in a concentric flower-like pattern, which is characteristic of *Mortierella*. Pure isolates were obtained after two successive rounds of subculturing onto PDA. We obtained seven different *Mortierella* isolates based on differences in their colony morphology. The identities of these isolates were determined by amplifying the full length internal transcribed spacer region (ITS) using the ITS1 and ITS4 primer pair [35] and Sanger sequencing.

### 2.4 Genomic DNA extraction and sequencing of *Mortierella* isolates

Total nucleic acid of *Mortierella* was extracted using the protocol from [36]. In brief, fungal tissues were harvested from PDA and powdered using liquid nitrogen and pre-chilled mortar and pestle. Preheated SDS lysis buffer containing RNaseA was added to the tissue powder. The lysate was homogenised by inversion, then incubated at 70°C for 30 min and inverted every 5 min. Contaminants were precipitated by adding 5 M potassium acetate and incubated on ice for 10 min. Debris was pelleted by centrifugation at 10,000 *x rcf* for 10 min. Nucleic acid in the aqueous phase were transferred to new tube and recovered by adding previously prepared magnetic Serapure beads and incubated at room temperature with gentle agitation for 15 min. Beads were pelleted by centrifugation at 10,000 *x rcf* for 3 min. Supernatant was decanted with care, then pelleted and washed with 70% ethanol twice. Beads were air-dried for 5 min and DNA eluted with 10 mM Tris-Cl pH 8 buffer preheated to 50°C. Libraries were constructed and paired end sequenced with Illumina HiSeq 2500 at Next-Generation Sequencing High Throughput Genomics Core at Biodiversity Research Center in Academia Sinica. Long-read sequencing of *Mortierella* was conducted using the Oxford Nanopore sequencing platform. A sequencing library was constructed using a Ligation Sequencing Kit (SQK-LSK109; Oxford Nanopore). Sequencing was carried out using R9.4 flowcells (FLO-MIN106) on a MinION (Mk1B) or GridION (Mk1) sequencer.

### 2.5 Transcriptome sequencing of *Mortierella* isolates

Approximately 50 mg of fungal tissue were transferred into a 2 mL screw-cap tube containing ceramic beads. The tubes were snap-frozen in liquid nitrogen for 5 min. Tissues were homogenised using PowerLyzer 24 set at 3,000 rpm for 20 sec. Tubes were removed from the homogeniser, temporarily allowing the contents to return to a liquid state, and snap-frozen again. The homogenisation process was repeated three times until a powder was obtained. The remainder of the extraction protocol was carried out using TRIzol (Cat. #15596026; Invitrogen) as instructed by the manufacturer. RNA sequencing libraries were prepared using Illumina poly-A stranded RNA library preparation kit and sequenced using Illumina HiSeq 2500 paired-end 2 × 150bp in rapid mode.

### 2.6 Construction and sequencing of amplicon libraries

Amplicon libraries were constructed by amplifying the ITS2 region using the protocol described by [11]. The DNA concentrations were quantitated using NanoDrop™ 1000 (ThermoFisher Scientific, Massachusetts, US). Amplicons were normalised using SequalPrep Normalization Plate Kit (ThermoFisher Scientific, Massachusetts, US). Normalised amplicons were pooled at equal volumes into a single tube, concentrated using AMPure XP beads (Beckman Coulter, California, US) at a 1:1 ratio and constructed into sequencing libraries using TruSeq DNA Preparation Kit (Illumina Inc, California, US). Sequencing was carried out using Illumina MiSeq paired-end 2 × 300 bp and performed by the High-Throughput Sequencing Core Facility in Biodiversity Research Center in Academia Sinica.

### 2.7 Amplicon sequence read pre-processing

Sequencing data generated from ITS libraries were demultiplexed using *sabre* (ver. 0.9). Reads were trimmed of primer sequences. Paired reads were merged and quality filtered using USEARCH v11.0.0 with parameters suggested by the UPARSE pipeline [37]. was used. Filtered sequence reads were removed of singleton and chimeric sequences and clustered into zero-radius operational taxonomic units (zOTUs) using the UNOISE3 algorithm [38]. Taxonomy was determined using the SINTAX algorithm [39] against the RDP database (v16). A zOTU table was generated using the *usearch_global* function.

### 2.8 Data analysis

Sample preparation variations were assessed by sequencing the Matsu archipelagos (MT) and Green Island (GI) in triplicate and quadruplet, respectively. Non-metric dimensional scaling (NMDS) suggested minor variation between technical replicates (**Supplementary Fig. 1**). Replicates were merged by calculating the mean of zOTU counts between replicates. 72.4% and 15.8% of the zOTUs were classified at the phylum and genus levels. Unclassified zOTU sequences were identified by BLAST against the ITS_RefSeq NCBI database [40] and non-fungal zOTUs were removed.

Outputs of the UPARSE pipeline were imported into the R environment using RStudio. Data were analysed using the *phyloseq* package (ver. 1.34) and diversity analyses were conducted with the *vegan* package (ver. 2.5-7). The *ordiR2step* function was used to perform the Akaike information criterion (AIC) with a forward selection model on the edaphic and climatic variables and fungal profiles. The statistical significance of the differences in diversity was calculated using analysis of variance (ANOVA), Tukey Honest Significant Difference test, and permutational multivariate analysis of variance (*adonis*) when appropriate. The intersecting zOTUs from different islands were calculated using *UpSetR* (ver. 1.4) and visualised with *ComplexHeatmap* (ver. 2.4.3) packages. A co-occurrence network was constructed using *fastspar* (ver. 1.0), a C++ version of the SparCC algorithm [41,42]. Network statistics were calculated using the built-in plugin ‘Analyze Network’ in Cytoscape (ver. 3.8.1). Node strength was calculated by averaging the absolute value of connected edge-weight per node. Modules in the biological network were determined using WGCNA [43]. The zOTUs were characterised into peripherals, connectors modules hubs and network hubs using within-module degree (z score) and among-module connectivity (c score) thresholds as previously described [44].

### 2.9 Assembly, annotation and phylogenomic analyses of *Mortierella* isolates

Raw sequence data from nanopore sequencing were basecalled using guppy (ver. 3.3.3). Read error correction was conducted using Canu (ver. 1.9) and *de novo* assembly was performed with Flye (ver. 2.5). The consensus sequence of the assembly was corrected with four rounds of Racon (ver. 1.4.6) and then once using Medaka (ver. 0.11.0). Low-quality Illumina bases were trimmed using Trimmomatic (ver. 0.36), and genome assembly was polished with Illumina sequencing data using five rounds of Pilon (ver1.22).

RNAseq reads of each *Mortierella* species were aligned to corresponding assemblies using STAR (ver. 2.7.2d [45]) and reference assembled using Stringtie (ver. 2.1.1 [46]) and Trinity (ver. 2.9.1 [47]). These transcripts were filtered and picked using MIKADO (ver. 2.0rc6 [48]). The passed transcripts were then used as hints and underwent an initial round of annotation using BRAKER2 (ver. 2.1.5[49]). The BRAKER2 predictions, protein sequences from uniport-fungi, *Rhizophagus irregularis* DAOM197198 and *Mortierella elongata* AG-77 from JGI, and RNAseq-assembled transcripts were used as input for MAKER2 (ver. 3.01.03 [50]) annotation pipeline to generate a final set of gene model predictions. BUSCO was run on the proteomes of seven *Mortierella* isolates (ver. 5.0.0 using fungi_odb10 and mucoromycota_odb10 database, [51]).

ITS phylogenetic tree was constructed using the 7 *Mortierella* isolates and 298 sequences of *Mortierella* strains [52]. Full-length ITS sequences of the 298 strains were retrieved from NCBI using Batch Entrez tool using accession number listed in the publication [52]. ITS sequence of soil isolates from this study was obtained using Sanger sequencing with ITS1 and ITS4 primer pairs [35]. Sequences were aligned using MAFFT (ver. 7.471) with options *--maxiterate 1000 --localpair*. Alignments were trimmed using TrimAL (ver. 1.2) with option *--automated1*. Phylogenetic tree was calculated from trimmed sequences using IQ-Tree with the options *--MF --bb 1000*, the best model determined was TIM2+F+R10.

Orthogroups of seven *Mortierella* isolates and 27 representative species (**Supplementary Table 1**) were identified using OrthoFinder (ver. 2.27, [53]). A maximum likelihood phylogeny was inferred using an concatenated alignment of single copy orthologs using FastTree (ver. 2.1.10, [54]) with 1000 bootstrap and options *-gamma -lg*.

### 2.10 Construction of custom fungal database for PICRUSt2

PICRUSt2 contains a built-in fungal database for functional inference based on ITS sequencing data. Instructions for using a non-default database and generating custom databases are detailed on the PICRUSt2 GitHub page (https://github.com/picrust/picrust2). Briefly, ITS sequences of *Mortierella* isolates were determined by Sanger sequencing with the ITS1 and ITS4 primer pair [35]. ITS regions of *Mortierella* genomes from the JGI database were identified by aligning against the *Mortierella* ITS sequence from the NCBI database using BLAST. The list of genomes incorporated are listed in **Supplementary Table 1**. A multiple-sequence alignment file containing newly determined ITS sequences and the existing sequences from PICRUSt2 were generated using MAFFT (v7.471). A phylogenetic tree file and a hidden-Markov model file were calculated from the new alignment using IQ-TREE (ver. 1.6.12) and *hmmbuild* (ver. 3.1b2), respectively. A model file was generated using RaxML (ver. 8.2.11, [55]). ITS copy number in the *Mortierella* genomes was determined by aligning against the NCBI ITS reference database using BLAST. Enzyme commission (EC) counts of the *Mortierella* genomes were extracted and tabulated from GFF files of *Mortierella* genomes. KEGG orthologues (KO) were inferred from EC number using the *KEGGREST* package (ver. 1.28.0) in R. The new ITS copy number, EC counts, and KO counts were appended to their respectively PICRUSt2 files.

## 3. Results

### 3.1. Data statistics

DNA samples of soils from nine Taiwan offshore islands (**Fig. 1**) were purified and characterised by sequencing the fungal ITS2 region using the Illumina MiSeq platform. A total of 3,711,303 reads with an average of 78,964 reads for each of 47 samples were used for downstream analysis. Deduplicated reads were denoised and decontaminated into 8,528 zero-radius operational taxonomic units (zOTUs) [38] with an average of 1,257 zOTUs per sample. A total of 94.8% and 92.0% of zOTUs were classified successfully at the phylum and genus level, respectively.

### 3.2. Comparison of soil fungal diversities between islands

An overview of fungal alpha diversity indices showed a high Chao1 index and low effective number of species (ENS) for all sampled islands (**Fig. 2**), with no strong correlation to edaphic data in either index (**Supplementary Fig. 2**). Higher fungal richness was observed in larger islands (**Supplementary Table 2;** MT**-**BG, MT-NG, MT-DJ, MT-SJ, OI and GI median Chao1 = 2,035.0, 1,719.4, 1,809.3, 1,910.2, 1,323.2 and 1,850.7, respectively), and small islets such as Dongyin (MT-DY), Dongsha (DS) and Taiping (TP) showed low fungal richness (MT-DY, DS and TP median Chao1 = 864.0, 976.4 and 373.7, respectively). In contrast to the variation in species richness, all islands were found to have similar ENS, except GI (**Supplementary Table 2**; all islands – mean ENS = 53.4, GI – mean ENS = 142.0), which suggests that island size impacted total fungal richness but not the number of representative taxa in the soil.

**Fig. 2.**
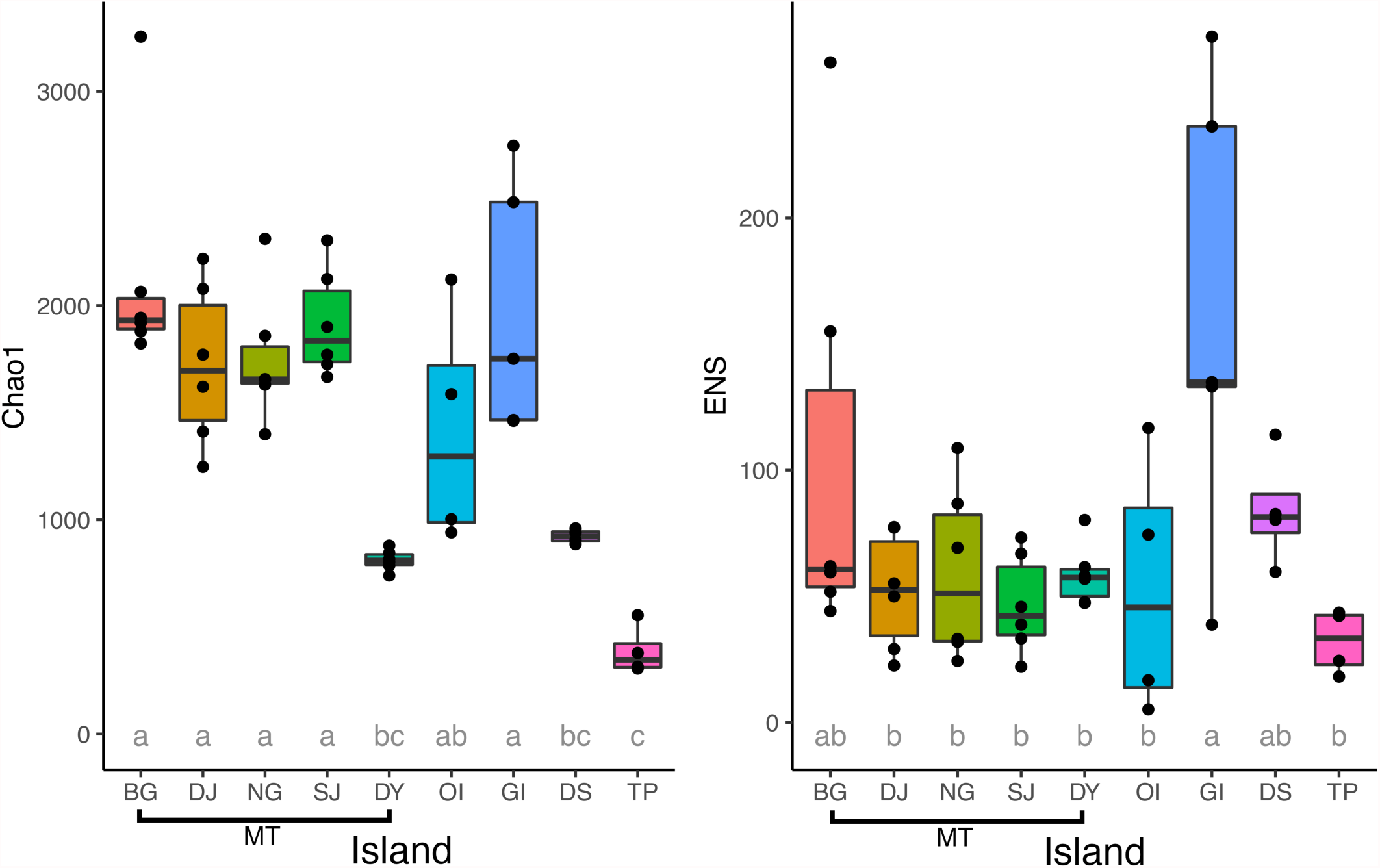
Fungal alpha diversity indices of the offshore islands. Richness and evenness are represented by Chao1 and effective number of species (ENS), respectively. Pairwise significant differences in statistical means between islands were calculated using ANOVA and Tukey HSD and denoted using letters. E.g. islands labelled *a* are not significantly different to island with the same letter. Islands labelled *ab* indicate they are not statistically significant to neither group *a* or *b*.

The species-area relationship on these nine islands revealed that island size is positively correlated to alpha and gamma diversities (**Fig. 3a**), but not beta diversity, suggesting an area *per se* effect. Distance-decay relationship analysis demonstrated that the mycobiome similarities between islands were inversely correlated with geographical distance (**Fig. 3b; left panel**). In comparison, intra-island mycobiome dissimilarities demonstrated a weak positive or no correlation with distance **(Supplementary Fig. 3)**, suggesting within an island the likelihood of species migration and colonisation remained high, thus reducing differences between communities. Independent analyses of the abundant (>1% relative abundance) and rare (<1% relative abundance) fungal taxa exhibited the same distance-decay trend in their community dissimilarities **(Fig. 3b; middle and right panels)**. However, rare taxa showed a higher level of dissimilarity (Bray-Curtis > 0.6) at a close distance than the abundant taxa suggesting that rare taxa are specialists in the environment, shaped by islands’ niches and remained relatively dissimilar regardless of distance.

**Fig 3.**
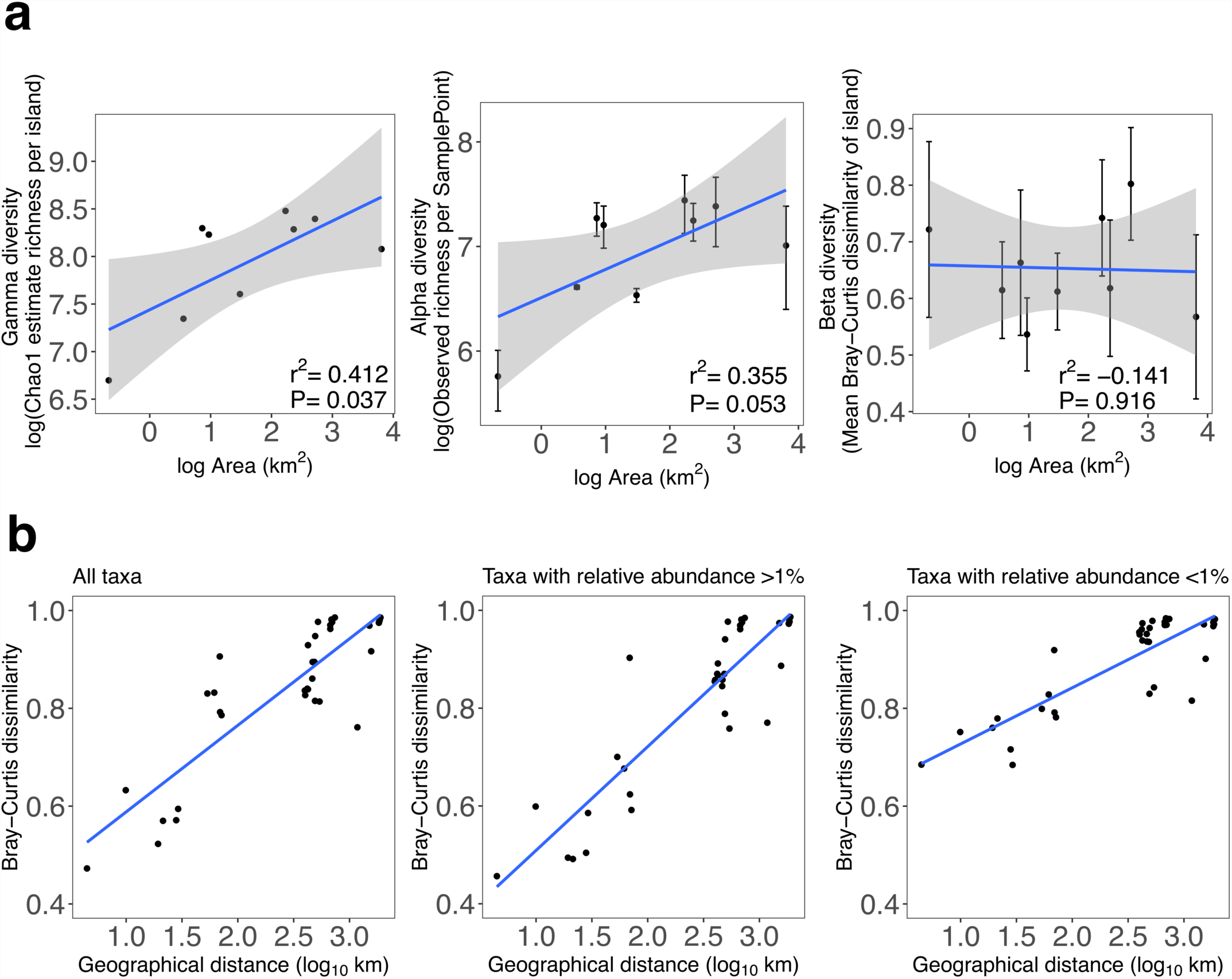
Species-area and distance-decay relationship of Taiwan’s offshore islands. **a**. Offshore island species-area relationship. Alpha diversity was calculated from the observed zOTU richness for each sample point. Averages represent the island and standard deviation from the mean is indicated with error bars. Beta diversity was calculated from the average Bray-Curtis distance within each island. Gamma diversity is represented by the Chao1 estimator of the island. **b**. Distance-decay relationship across Taiwan’s offshore islands. Each point represents a pairwise community difference between islands. Linear regression was used to calculate the line of best fit.

The mycobiome structures between islands were driven by island soil type, as the samples grouped in the NMDS space according to their island parent rock material (**Fig. 4a**). For example, soil mycobiomes of MT (granite) formed an independent cluster. Interestingly, mycobiomes of GI (andesite) and DS (coral) were found to be alike despite the different soil types and geographically distant. Overall, PERMANOVA analysis of fungal profiles between all islands showed significant differences between NG and MT-DY only (P=0.036), suggesting that the communities were similar across all islands. We collected 17 edaphic and climatic metadata, and 10 of the 17 variables were found to be correlated with the soil mycobiome (**Supplementary Table 3**). Fungal decomposers (Mortierellomycota and Mucoromycota) and catabolic enzyme activities (cellulase, xylanase and β-glucosaminidase) exhibited a positive correlation with MT mycobiomes. The soil variables MBC, MBN, pH and the abundance of Ascomycota were correlated with the mycobiome of GI, DS and TP (**Fig. 4b**). The soil samples from OI did not correlate with any metadata or specific fungal phyla.

**Fig. 4.**
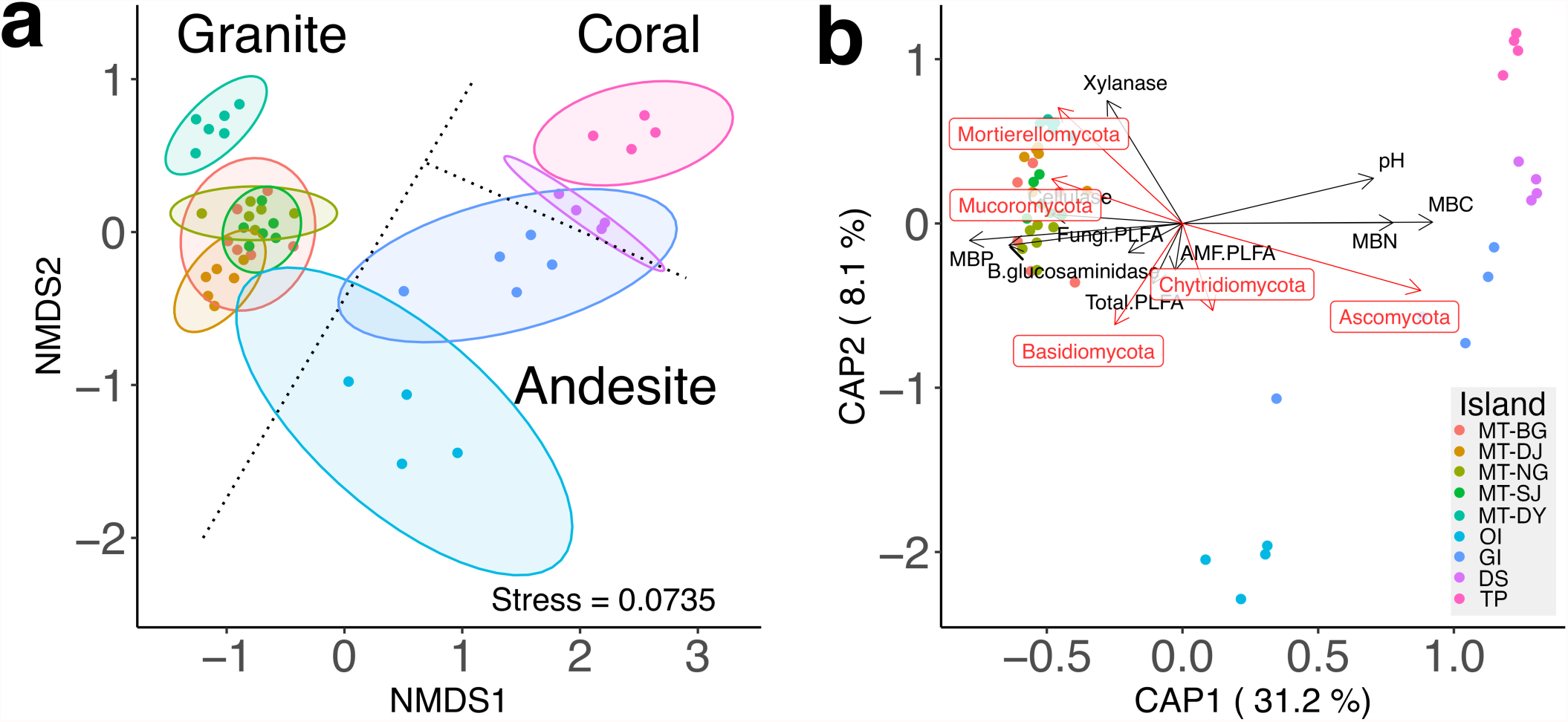
Ordination analysis of the offshore island soil mycobiomes. **a**. Non-metric dimensional scaling (NMDS) of fungal profiles. **b**. Biplot diagram showing the constrained ordination (distance-based redundancy analysis) using edaphic data and the abundance data from the five most dominant phyla. Points represent sampling sites and colours denote different islands.

### 3.3 Comparison of taxonomic profile and fungal abundance revealed dominance of *Mortierella* in the soil

The overview of the fungal composition profiles showed a high abundance of either Mortierellomycota (MT-BG, MT-NG, MT_DJ, MT-SJ and OI) or Ascomycota (GI, DS and TI; **Fig. 5a**). Interestingly, OI was found to have a high proportion of Mortierellomycota (mean abundance = 60.3%), albeit distant from MT and with different soil types. We found that the genus *Mortierella* alone comprised 11–65% of the total fungal relative abundance. Strikingly, nearly half of the total fungal abundance was dominated by only five *Mortierella* zOTUs on four of the nine islands (**Fig. 5b**).

**Fig. 5.**
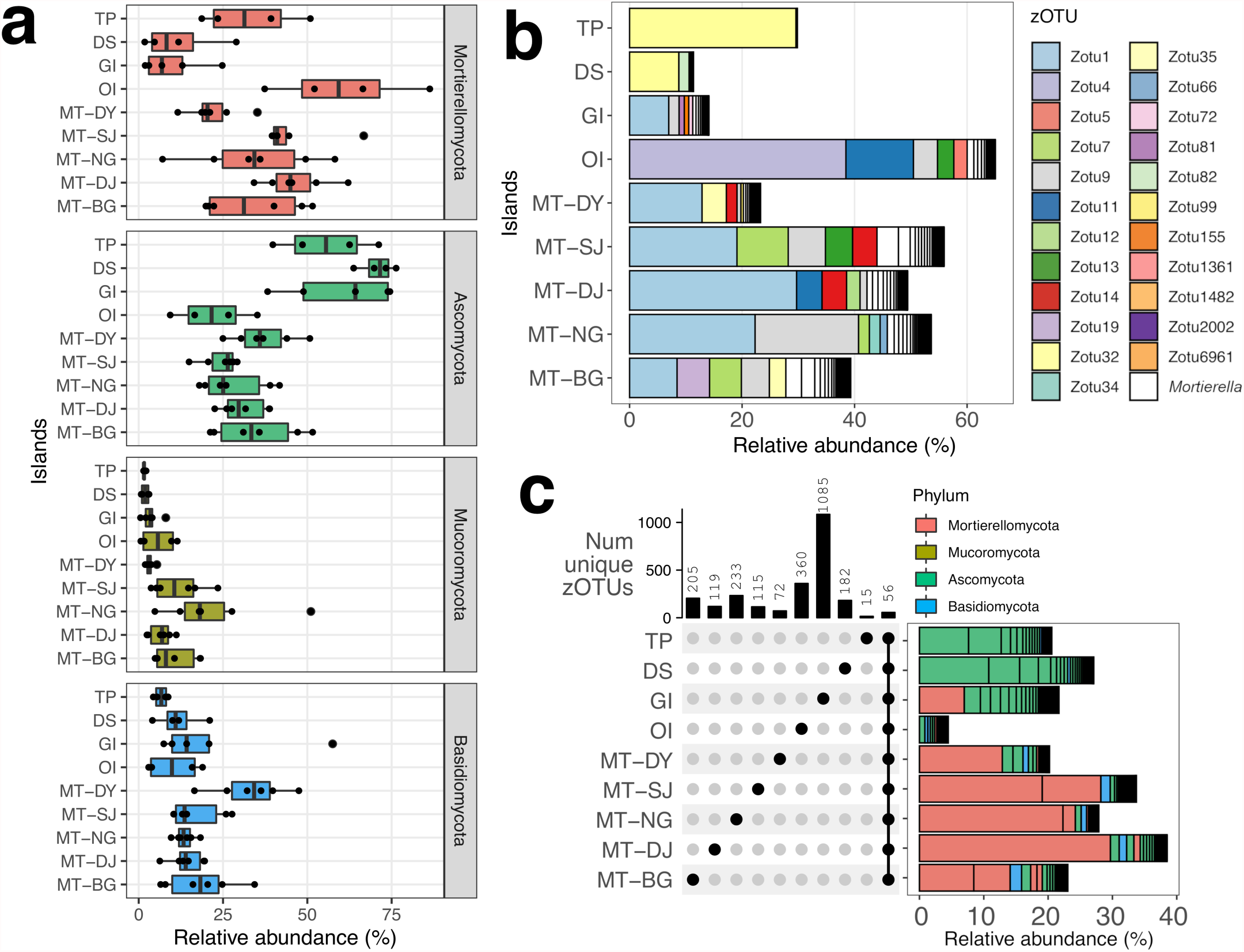
Relative abundance and composition of the offshore islands soil mycobiome. **a**. Summary of the relative abundance of the four most abundant phyla on the nine offshore islands. Points represent the sample sites. **b**. Relative abundance of the genus *Mortierella* on each island; the five most abundant *Mortierella* zOTUs from each island are denoted by different colours. **c**. Upset diagram showing the number of overlapping and unique zOTUs from each island. Adjoined bar plot shows the relative abundance of the 56 cosmopolitan zOTUs on each island; phyla are denoted by different colours.

Fifty-six zOTUs were found across all nine islands (hereafter termed cosmopolitan zOTUs), contributing on average 24.12% of the samples’ relative abundances (maximum, minimum and median relative abundance contributed = 38.5%, 4.48% and 23.06%, respectively). Furthermore, the abundance composition profiles contributed by the cosmopolitan zOTUs were similar to the dominant fungal phyla on each island, except for OI (**Fig. 5c, Supplementary Table 4**). While the cosmopolitan zOTUs represented the Mortierellomycota abundance in MT, they only contributed marginally to the relative abundance in OI (4.48%), indicating that unique *Mortierella* taxa were present.

### 3.4 Correlation network analysis revealed *Mortierella* as a keystone species in soil mycobiome

The putative importance of the dominant genus *Mortierella* and its relationship with other species were explored using network analysis. The number of interacting zOTUs and interaction strengths were represented by the zOTU degreeness and node strength, respectively. A similar degreeness was detected among different phyla on each island. Mortierellomycota and Mucoromycota exhibited higher median degreeness, albeit not significant except for MT-NG (**Fig. 6a;** ANOVA, Mortierellomycota-Mucoromycota P=0.011; Mortierellomycota-Ascomycota P=0.037; Mortierellomycota-Basidiomycota P=0.001; **Supplementary Table 5**). Mortierellomycota-dominant islands showed Mortierellomycota zOTUs with high node strength compared to other phyla (**Fig. 6a, Supplementary Table 6)**. zOTUs were next classified into peripheral, connector, module hub or hub nodes within a network. The majority of the zOTUs were defined as connector and peripheral nodes (**Fig. 6b**). In total, 26 network hubs and 19 module hubs were characterised from the nine islands (**Fig. 6c**). The seven of the 45 hub species (**Supplementary Table 7**; zOTU1 - *Mortierella*, zOTU7 - *Mortierella*, zOTU21 - *Clonostachys*, zOTU75 - *Trichoderma*, zOTU226 - *Clonostachys*, zOTU304 - *Staphylotrichum*, and zOTU333 - *Aspergillus*) were ubiquitous across islands; these genera are readily found in the environment as saprophytes [11]. *Mortierella* (zOTU1 and zOTU7) and *Trichoderma* (zOTU75) species were correlated with degradative enzyme activities (**Supplementary Fig. 4**). Together, these results suggest that *Mortierella* is an integral member of the soil community. It is highly connected to other species, hence classified as hub nodes, and shown by the high intra-network and inter-module connectivity.

**Fig 6.**
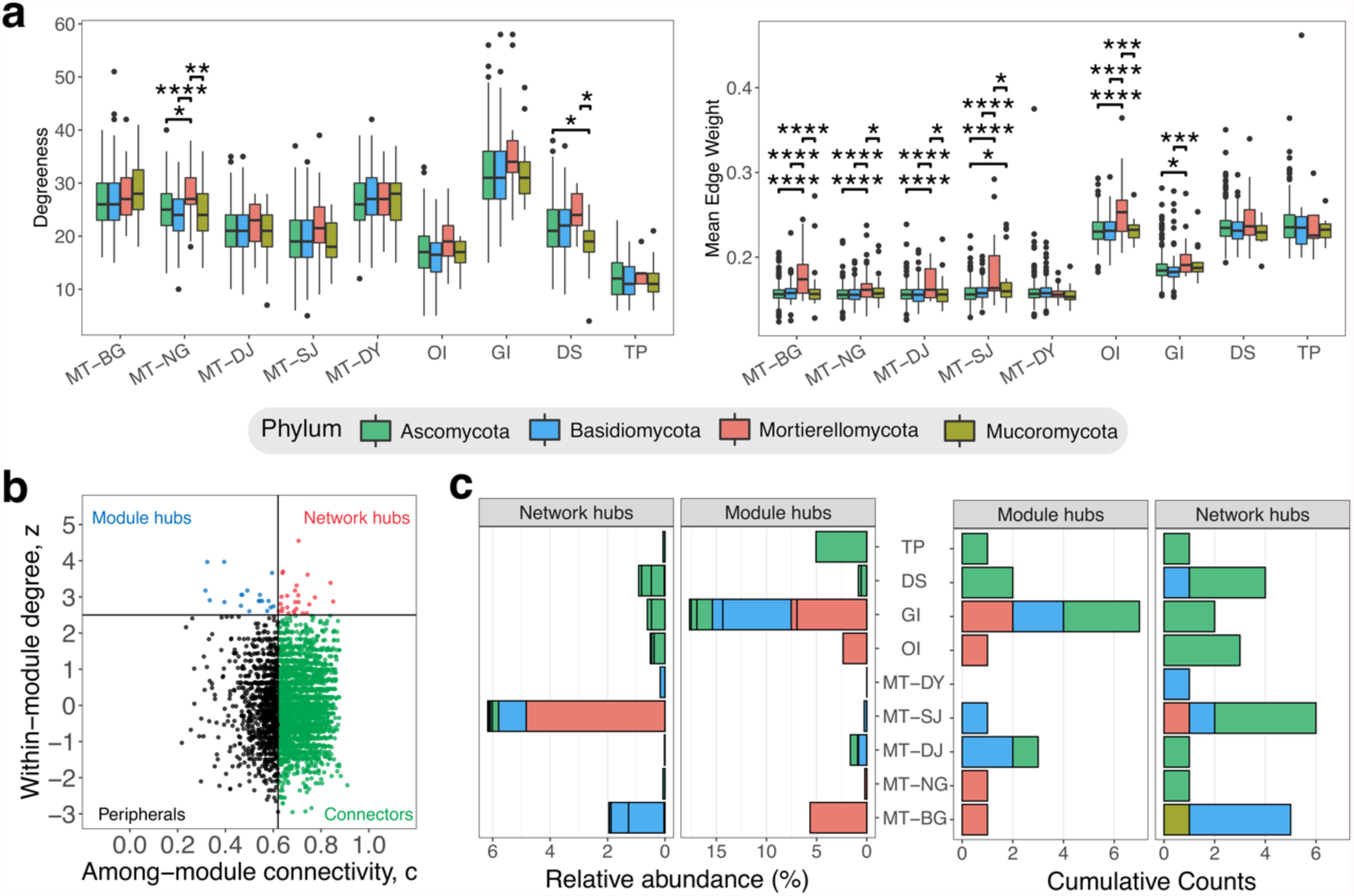
Correlation network statistics of the offshore islands soil mycobiome. **a**. Degreeness and mean edge weight (node strength) distribution of the four most dominant fungal phyla from correlation networks on each island. Significant differences between fungal phyla within the island are labelled with asterisks (P ⩽ 0.05 = *; P ⩽ 0.01 = **, P ⩽ 0.005 = ***; P ⩽ 0.001 = ****). **b**. Classification of zOTUs into hubs, connectors or peripherals nodes. **c**. Number of module and network hubs, and their respective relative abundance in each island.

### 3.5 *De novo* assemblies and phylogenomics of Taiwanese *Mortierella* isolates

Motivated by our results that *Mortierella* is an important genus in the soil fungal community, we isolated, sequenced and annotated seven *Mortierella* isolates (Methods). The assemblies range 34.5-55.5Mb in size from 313,449-1,313,866 Oxford Nanopore long reads which were subsequently polished with Illumina reads (average Nanopore read N50 = 20.8kb; contig N50 = 3.65Mb; **Supplementary Table 1**). These assemblies were highly contiguated compared to published *Mortierella* genomes, for example the new *M. elongata* SJ01-01assembly has a N90 of 1.68Mb compared to *M. elongata* NVP64 with N90 of 1.15Mb. Using the available *Mortierella* and transcriptome sequencing from mycelium, we annotated 8,389-13,336 gene models using the MAKER2 pipeline (**Supplementary Table 1**). Assessment the completeness of annotation using BUSCO (benchmarking universal single-copy orthologs) suggest that they are 95.0-98.4% complete, which is comparable to available proteomes in these species (**Supplementary Table 1)**. Finally, we attempted to place these species phylogenetically by constructing a phylogeny either from i) ITS (**Supplementary Figure 5**) or ii) 528 single copy orthologues (**Fig. 7**). Based on previously described classification system [56], we classified both isolate SJ01-01 and BG05-11 as *M. elongata*, NG01-01 as *M. minutissima*. BG05-04, and SJ01-07 were placed in the Gamsii clade sister to *M. elongata*. Two isolates NG01-10 and NG01-12 were grouped with *M. wolfii* and *M. alpina* in the ITS phylogeny, respectively, but were singly placed without any species in the species phylogeny, suggesting that they were the first assembly for these species (**Fig. 7**).

**Figure 7.**
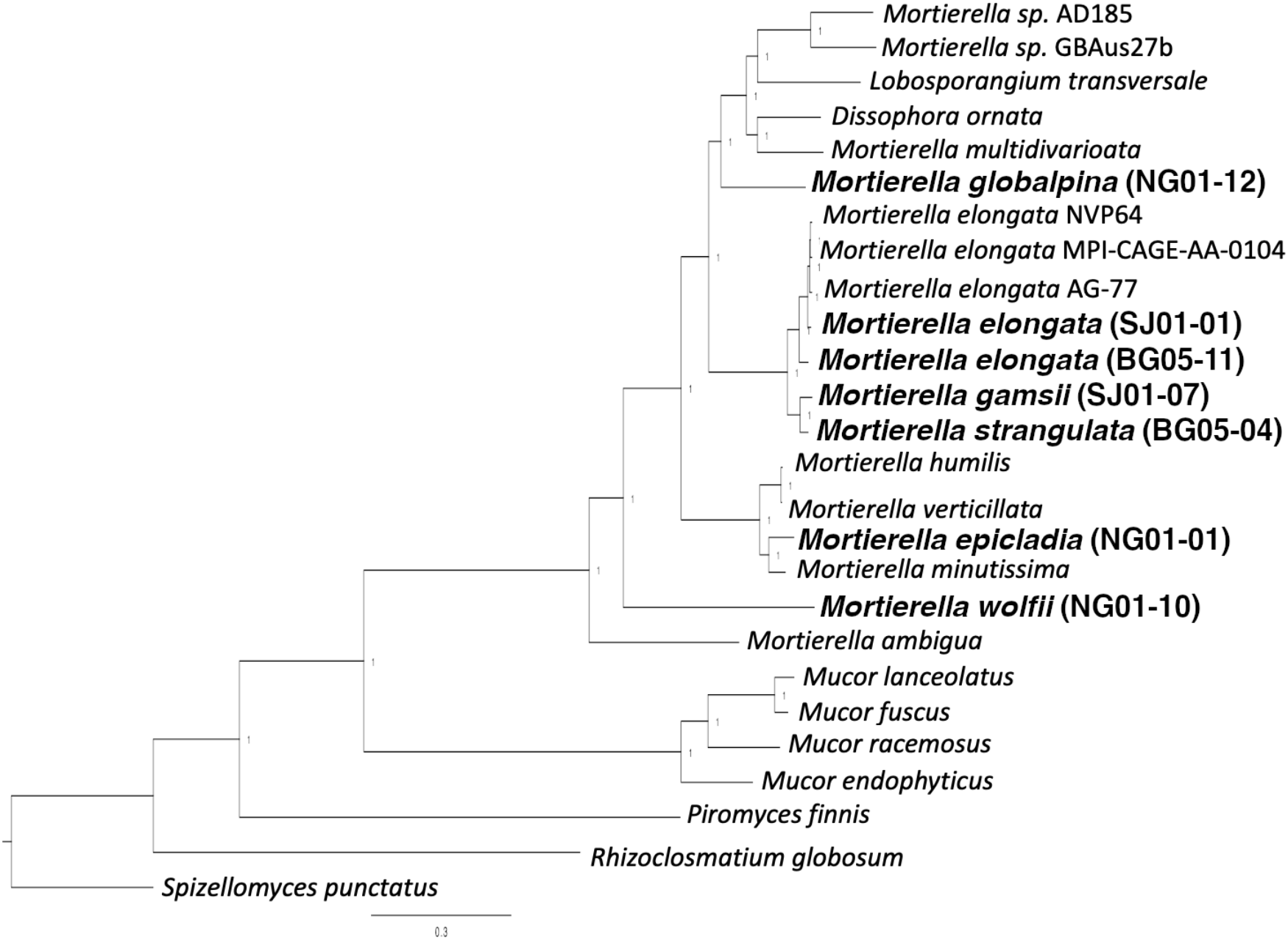
Phylogenetic tree of *Mortierella* genomes based on 528 single copy orthologs. Text in bracket denote isolate ID. Bald letter indicate species made available in this study.

### 3.6 *Mortierella*-annotated PICRUSt2 database improves functional predictions for the surface soil mycobiome

We incorporated genomic data from the *Mortierella* genomes, which increased the putative relative abundance coverage of PICRUSt2 by 62.91% and 31.41% at the genus and phylum level, respectively, compared to covering 8.28% (141 zOTUs) and 66.34% (8,153 zOTUs) at the genus and phylum level, respectively (**Fig. 8a**). The ordination of KO profiles indicated that *Mortierella* abundance is a significant variable (*adonis*, P=0.01) regardless of whether or not *Mortierella* genomes are added. However, visual inspection of the improved database showed a tighter grouping for islands with high *Mortierella* abundance and a high percent variation explained on the first principle component (Original PC1 vs improved PC1; 41% vs 58%; **Fig. 8b**).

**Fig. 8.**
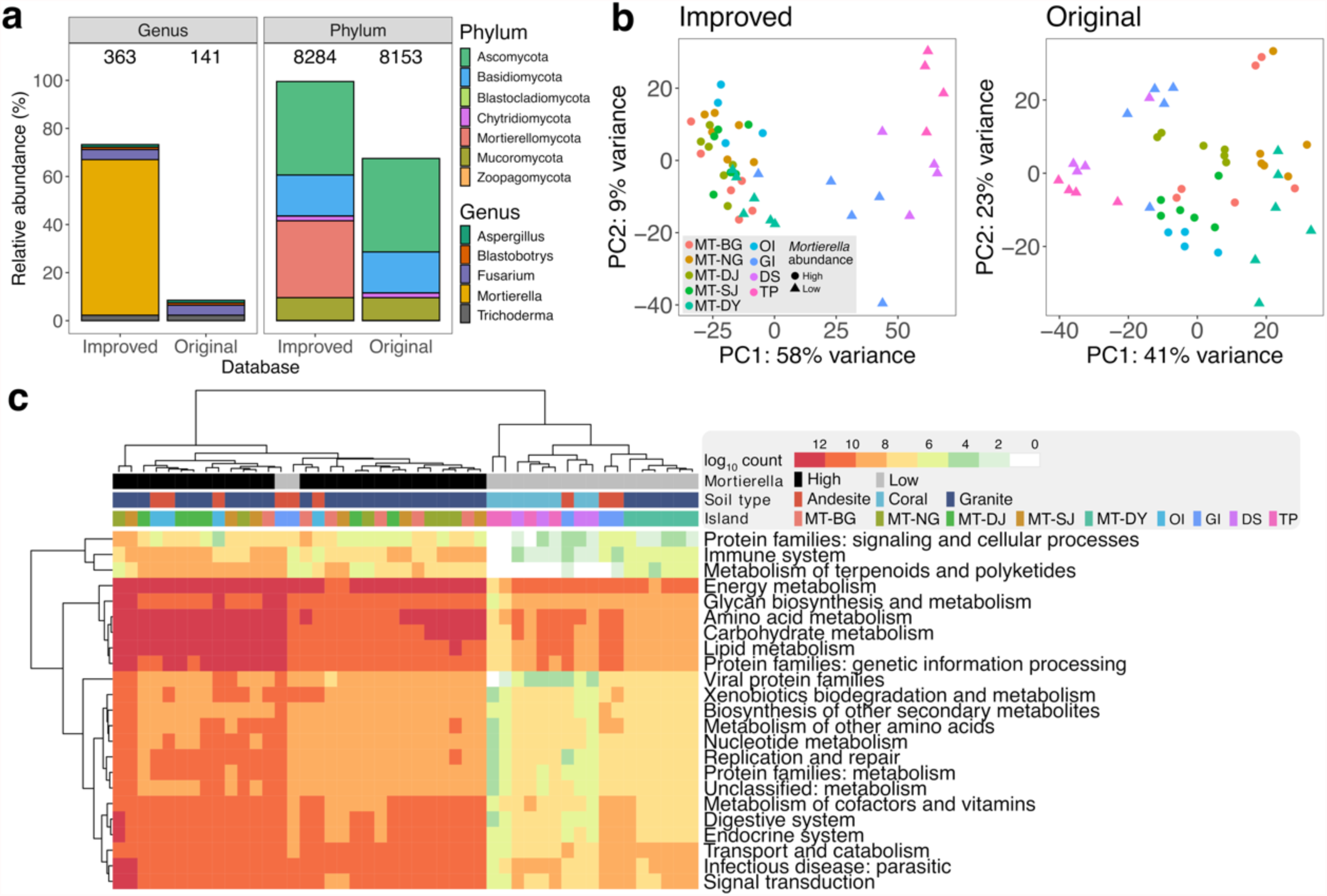
**a**. Estimated relative abundance coverage of the soil mycobiome at the genus and phylum levels comparing results from the improved and original PICRUSt2 pipelines. Number of zOTU covered are shown at the top of the bar plot. **b**. Principal coordinate analysis of the predicted KO profiles. Each point represents a sample site. Colours and shapes denote a different island and *Mortierella* abundance, respectively. **c**. Heatmap showing differentially abundant KOs predicted using the improved PICRUSt2 database grouped based on KEGG BRITE hierarchy level 2.

To highlight the difference in prediction results between the improved and original databases, differentially-abundant KOs and pathways were determined from DESeq2 analyses. No notable differences were observed in the number of differential pathways predicted from both databases; however, the improved database predicted almost 10 times more differential KOs than the original database (**Supplementary Table 8**. original vs improved; 25 vs 243 and 19 vs 133 for increased and decrease KOs, respectively). The heatmap indicated that differential KOs clustered according to *Mortierella* abundance, and not with the island *per se* or soil type (**Fig. 8c**). KOs with the most significant differential increase were detected in Energy metabolism, Glycan biosynthesis and metabolism, Amino acid metabolism, Carbohydrate metabolism, Lipid metabolism and Protein families: genetic information processing (**Fig. 8c**). Predictions made using the original database did not show significant changes in these categories (**Supplementary Fig. 6**). In the gene-level analysis, *Mortierella*’s role as a decomposer in soil was further highlighted by the increase we observed in the number of predicted carbohydrate-active enzymes (CAZy)—e.g., the differential increase in hexosaminidase (HEX), part of the glycosyl hydrolase 20 family (GH20). These widely distributed genes catalyse glycosidic-linked N-acetylhexosamine residue cleavage in N-acetylglucosamine and N-acetylgalactosamine, which plays a vital role in chitin, and cell wall turnover [57,58] (**Supplementary Fig. 7)**. Malate dehydrogenase (*mdh*) has been reported to be expressed by filamentous fungi in association with the degradation of biopolymers [59] (**Supplementary Fig. 8**). The fatty acid production ability of *Mortierella* was also reflected by the significant increase in acetyl-CoA carboxylase, which controls the production of malonyl-CoA, a vital intermediate substrate, in fatty acid biosynthesis and degradation [60,61] (**Supplementary Fig. 9**). Other enzymes associated with fatty acid metabolism were predicted, such as the phospholipase A2 (PLA2) [62], UTP-glucose-1-phosphate uridylyltransferase (UGP3) and glycerol dehydrogenase (*golD*) [63] (**Supplementary Fig. 10**). Altogether, these results suggest our ability to correlate *Mortierella*’s role in the soil as a saprophyte [64] and free fatty acid producer [65] using improved functional prediction workflow.

## 4. Discussion

Amplicon sequencing remains a highly efficient and cost-effective method to obtain a holistic view of the niche species compositions and functions in microbial ecology. We have characterised the soil mycobiome from 141 samples across nine islands, highlighted *Mortierella* dominance as keystone species. The availability of the 17 new *Mortierella* genomes provided the genomic resource for the curation of PICRUSt2 database, which revealed differentially functional genes associated with ecological roles of *Mortierella* on islands where it is dominant, otherwise could not be detected from default database. Our approach demonstrated the importance of providing genomic data from key taxa in amplicon gene prediction studies to better reflect the mycobiome functional roles. The improved database has revealed significant differences at the gene levels (KO) but not at the pathway level. This further emphasis the functional redundancy phenomenon in microbial [66] and fungal communities [67]. This has provided initial insights into soil mycobiome functions in association with biogeography patterns, where we observed alike functions from mycobiome in independent islands with significant compositional differences. In addition, it is worth noting that Ascomycota, which consisted 4,955 of the zOTU covered (38.2% of total zOTUs), remained poorly represented at the genus level even for this relatively well-studied phylum. This calls for attentions to the continual need to expand the existing fungal genome database to ascertain details about fungal functional relationship with respect to biogeography theories.

Intuitively, highly taxa abundance are associated with being a key member or having a significant impact on their niche [68,69]. Pervasiveness, persistence and being highly connected to other species are typical characteristics that enable keystone species to orchestrate community functions for adaptation. However, keystone species are not necessary the most dominant population in the community [70]. Core bacterial taxa in agricultural soil has been shown with a wide-range of relative abundance irrespective of their role in nutrient cycling [71]. The 56 cosmopolitan zOTUs are putative core soil taxa due to their pervasiveness. Our study has demonstrated that *Mortierella* species stood out from the rest of cosmopolitan zOTUs, possessing multiple keystone species attributes. *Mortierella’s* role as decomposers were reflected with high correlation to degradative enzyme activities. Interactions of *Mortierella* with other cosmopolitan zOTUs for community-level functions or temporal persistence of these putative core taxa warrants further investigation.

We demonstrated that insular fungal diversity is positively correlated with island area and distance, which congruent with the original island biogeography theory [72]. However, beta diversity remained equally dissimilar despite difference in island size. We hypothesised that colonisable space was not the limiting factor for fungal diversity unlike large and higher order organism such as plants and mammals. A lowered extinction rate and colonisation rate are associated with larger island; habitat and species diversity are more likely to be maintained, hence similar level of beta diversity. Despite the fact our sampling strategies focus on the region of the island away from human activities. We acknowledge that this resulted in islands not being extensively sampled. Thus, it is likely the increase in the island size did not truly reflect the increase in the variety of niches.

Our finding showed coral islands are coupled with lower fungal and plant diversity compared to other islands. Coral are highly porous; the low trace-nutrients and water-retention properties do not favour plant growth [73]. While plant being a significant determinant for fungal diversity [11], this translates to lowered soil fungal diversity as reported in other oceanic coral islands [74]. Microbiome study of Taiwan’s offshore island have highlighted the difference and important in soil type and the distinct microbiome structure observed between the Matsu archipelagos, GI and OI [33]. Soil nutrient dynamics are inherently linked with soil properties, which affect microbial growth physiology; therefore, it was expected for mycobiome to cluster according to rock type. A comprehensive edaphic data is also critical to explain putative role and presence of certain species. For example, MBP was positively correlated with mycobiome in granite islands. This correlation may be accounted by the high Mortierellomycota abundance, which are known for their role in soil phosphate solubilisation [75].

## 5. Conclusion

To conclude, the identification of incorporation of key fungal taxon to functional inference pipeline has significantly increased the amount of sequence abundance covered as well as revealed otherwise unpredicted differential KO associated with *Mortierella* metabolism. We believe the principle of this workflow can be tested on a perturbed system to better delineate the relationship between changes in microbial composition and pathway as well as biogeography relationship and overall functional changes in the mycobiome. The global effect in the continual discovery and curation of fungal genomes will undoubtedly aid *in silico* studies in the future.

## Acknowledgement

This research was funded by the Ministry of Science and Technology, Taiwan (Grant nos. 107-2313-B-001-003 and 107-2621-M-001-001). The funders had no role in the study’s design, data collection and analysis, decision to publish, or preparation of the manuscript. We thank members of Dr. Chih-Yu Chiu’s laboratory especially Pei-Yi Yu for assistance with field sampling and the collection of edaphic data. We thank the following for permission to use the following assemblies from JGI for species phylogeny and inferred metagenomics analyses: Gregory Bonito for *Mortierella* sp. AD185, *Mortierella* sp. GBAus27b, *Mortierella gamsii* AM1032, *Mortierella elongata* NVP64, *Mortierella humilis* PM_1414, *Mortierella minutissima* AD051 and *Mortierella ambigua* NRRL 28271, and *Mortierella wolfii* NRRL 6351, Joseph Spatafora for *Dissophora ornata* CBS347.77, *Mortierella multivaricata* RSA2152T, *Lobosporangium transversale* NRRL 3116, Rytas Vilgalys and Andrii Gryganskyi for *Mortierella elongata* AG-77.

## Authors contribution

I.J.T conceived and led the study. Y.F.L, I.J.T, Y.C.L, Y.J.L, E.H.C and C.Y.C carried out the sampling. Y.F.L, W.A.L and Y.J.L conducted the experiments. M.J.L and her team conducted experiments regarding high-throughput sequencing. W.A.L isolated the *Mortierella* strains and carried out the nanopore sequencing. H.H.L, Y.C.L, I.J.T carried out the sequence assembly and annotation. Y.F.L carried out the amplicon, correlation network and PICRUSt2 analysis. Y.F.L wrote the manuscript with inputs from I.J.T and C.Y.C.

## Data availability

Raw data, assemblies and annotation of the seven *Mortierella* isolates were deposited in the National Center for Biotechnology Information (accession no. PRJNA778874).

## Abbreviation

ITS: Internal transcribed spacer
zOTU: zero-radius operational taxonomic unit
KEGG: Kyoto Encyclopaedia of Genes and Genomes
KO: KEGG orthologues
PICRUSt: Phylogenetic investigation of communities by reconstruction of unobserved states
PDA: Potato dextrose agar
PBS: Phosphate buffered saline
PLFA: phospholipid-derived fatty acid
MBC: Microbial biomass carbon
MBP: Microbial biomass phosphorus
MBN: Microbial biomass nitrogen
AMF: Arbuscular mycorrhizal fungi
MMT: Mean monthly temperature
MMP: Mean monthly precipitation
MT: Matsu archipelagos
MT-BG: Beigan Island
MT-NG: Nangan Island
MT-DJ: Dongju Island
MT-SJ: Hsiju Island
MT-DY: Dongyin Islet
OI: Orchid Island
GI: Green Island
DS: Dongsha Islet
TP: Taiping Islet
NMDS: Non-metric dimensional scaling

