## Supplementary Info for "Genomes of keystone *Mortierella* species lead to better *in silico* prediction of soil mycobiome functions from Taiwan’s offshore islands"

### Supplementary Figures

**Supplementary Figure 1. NMDS plot of soil samples in Matsu Islands and Green Island**
Each of the soil samples were sequenced in triplicates. NMDS of mycobiome from replicate sample sequence of Matsu archipelagos (left) and Green Island (right) were plotted to visualise the data consistency between technical replicates. Colours denote different sample points on the island.

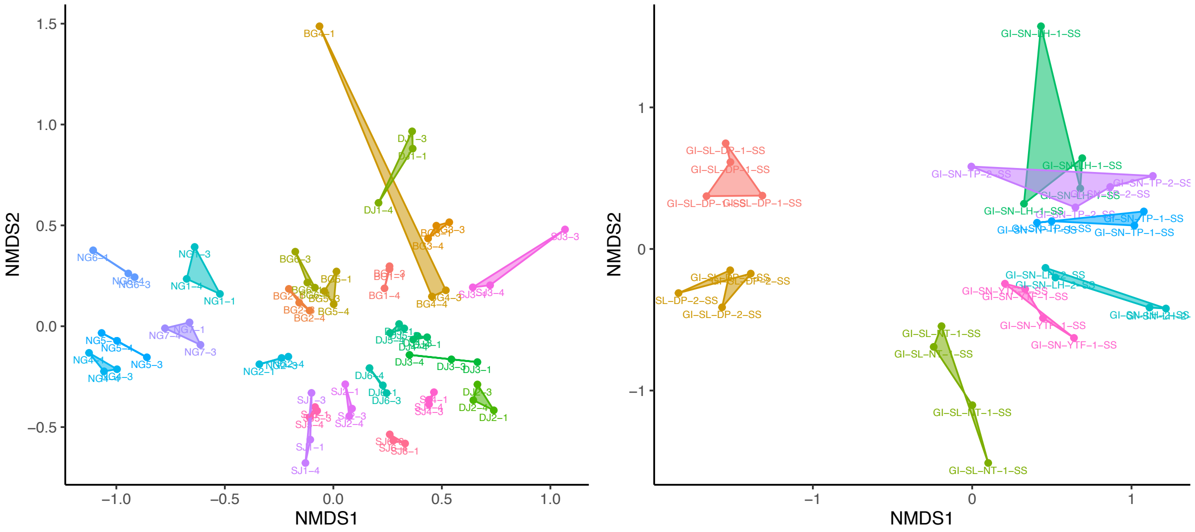

**Supplementary figure 2.** Correlation biotic and abiotic metadata against **a.** Chao1 diversity and **b.** effective number of species. Dot represent each sample point. The line of best fit indicates the correlation between the variable and alpha diversity index by linear regression.

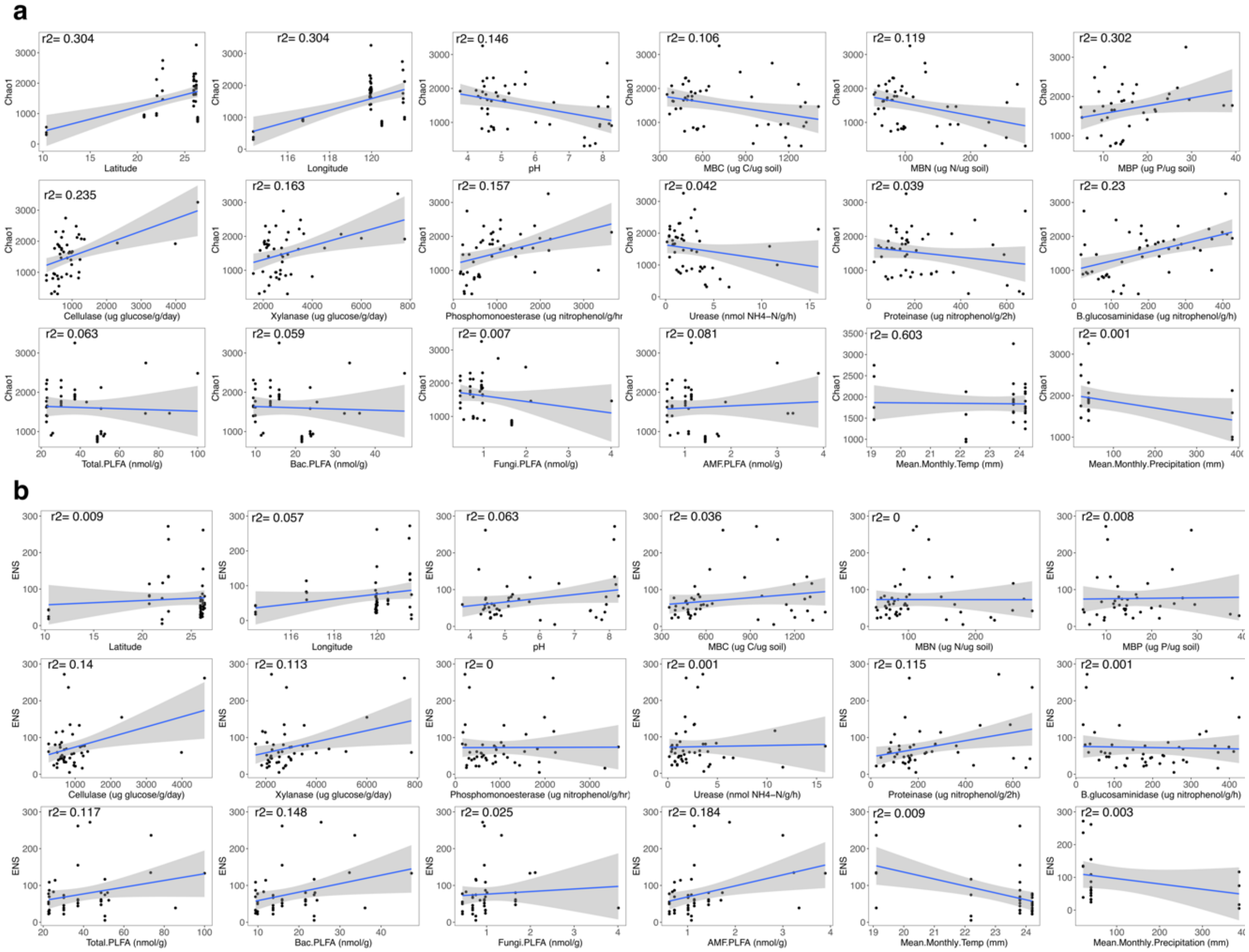

**Supplementary figure 3.** Community dissimilarity and distance relationship. Each point denotes the pairwise intra-island community dissimilarity between sample points.

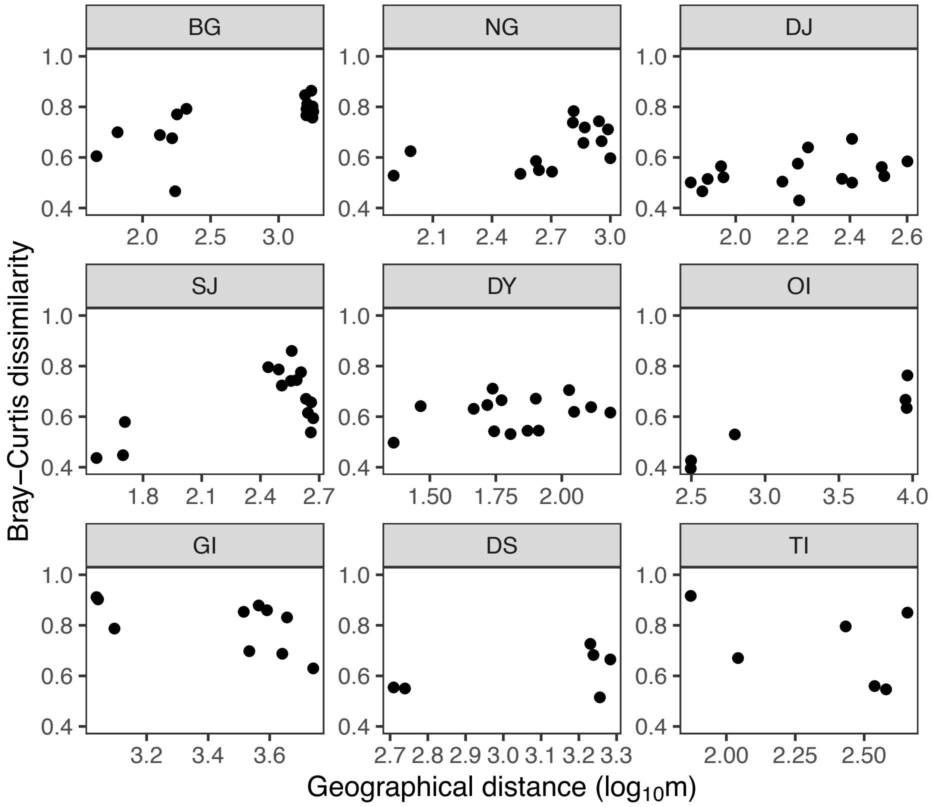


**Supplementary figure 4.** Biplot analysis of the island mycobiome with hub nodes abundance and edaphic metadata as explanatory variables.

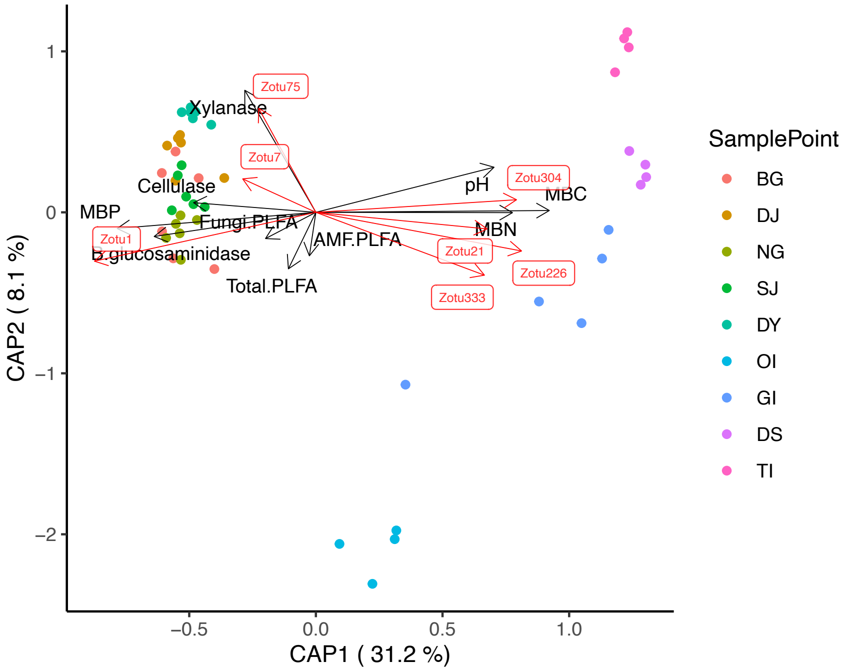

**Supplementary figure 5**. Phylogenetic tree of *Mortierella* genus constructed from 298 species obtained from NCBI database and the 7 isolates sequenced in this study. Phylogenetic classification of BG05-04 using ITS sequence showed to be placed distant from the Gamsii clade and appear to be an outgroup. However, aligning the ITS sequence of BG05-04 against the ITS reference database in NCBI revealed *M. gamsii* as the top hit (Percentage identity = 97.2%, E-value = 0, coverage = 87%).

Supplementary Figure 5 is in a separate pdf file.

**Supplementary figure 6.** Differentially abundant KO terms predicted using the original PICRUSt2 database grouped at BRITE hierarchy Level 1. Each row and column represent a KO term and soil sample, respectively. The heatmap represent $\log_{10}$ count predicted abundance of the relative KO terms. Soil sample characteristics (Island, soil type, *Mortierella* dominance) are shown by the top three rows. Heatmap was clustered using default setting of *hclust*. Subsequent supplementary figures (**Supplementary 7-10**) uses the same annotation format.

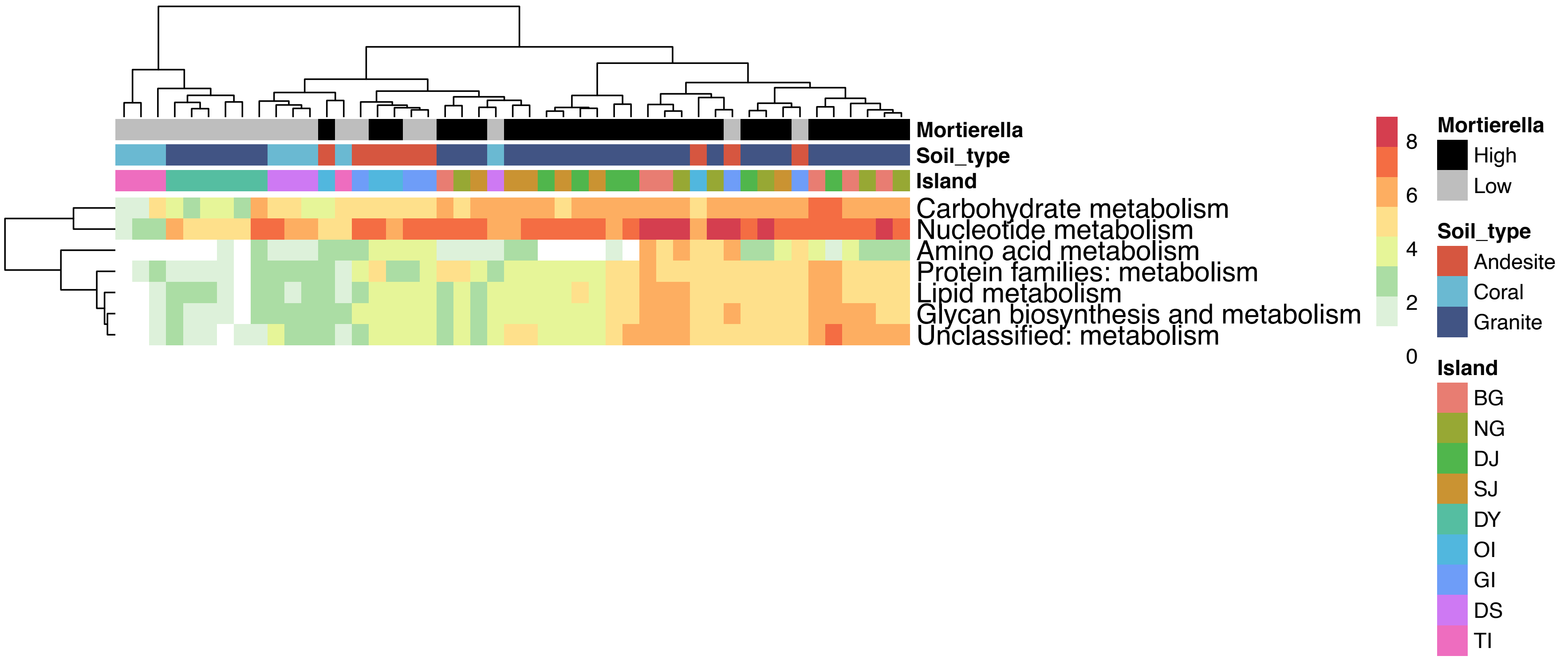

**Supplementary figure 7.** Differentially abundant glycan metabolism KO terms predicted using the improved PICRUSt2 database. Refer to Supplementary figure 6 for the annotation of this figure.

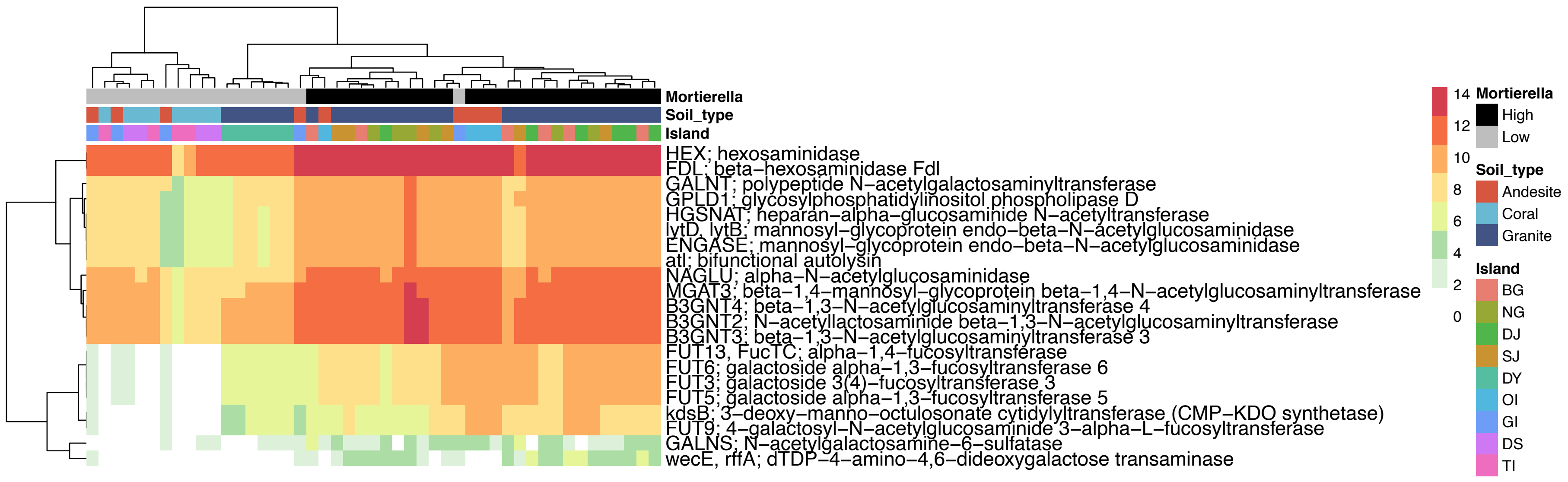

**Supplementary figure 8.** Differentially abundant carbohydrate metabolism KO terms predicted using the improved PICRUSt2 database. Refer to Supplementary figure 6 for the annotation of this figure.

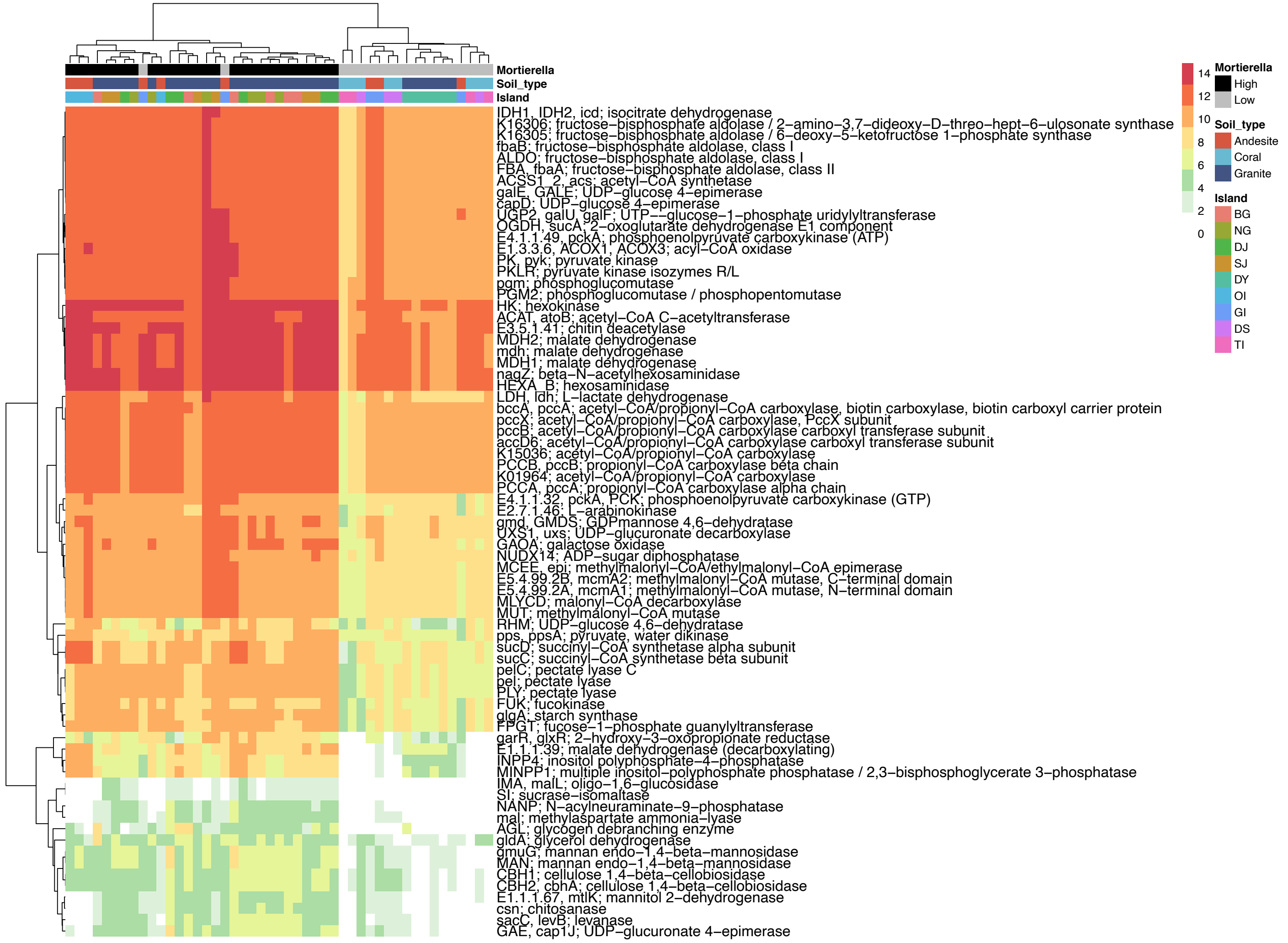

**Supplementary figure 9.** Differentially abundant energy metabolism KO terms predicted using the improved PICRUSt2 database. Refer to Supplementary figure 6 for the annotation of this figure.

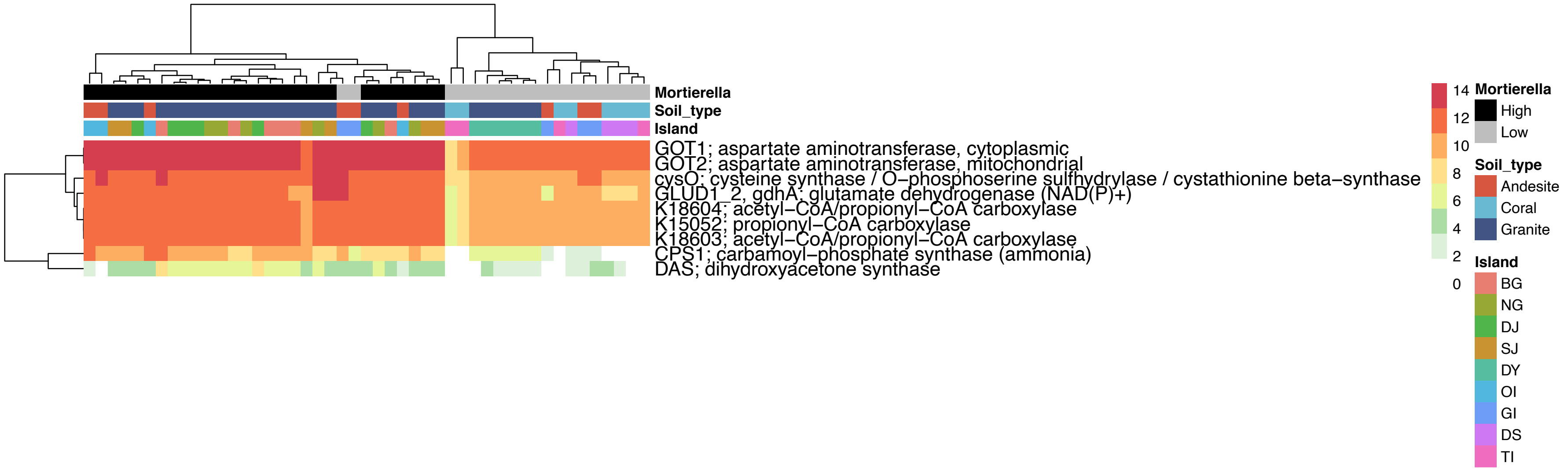

**Supplementary figure 10**. Differentially abundant lipid metabolism KO terms predicted using the improved PICRUSt2 database. Refer to Supplementary figure 6 for the annotation of this figure.

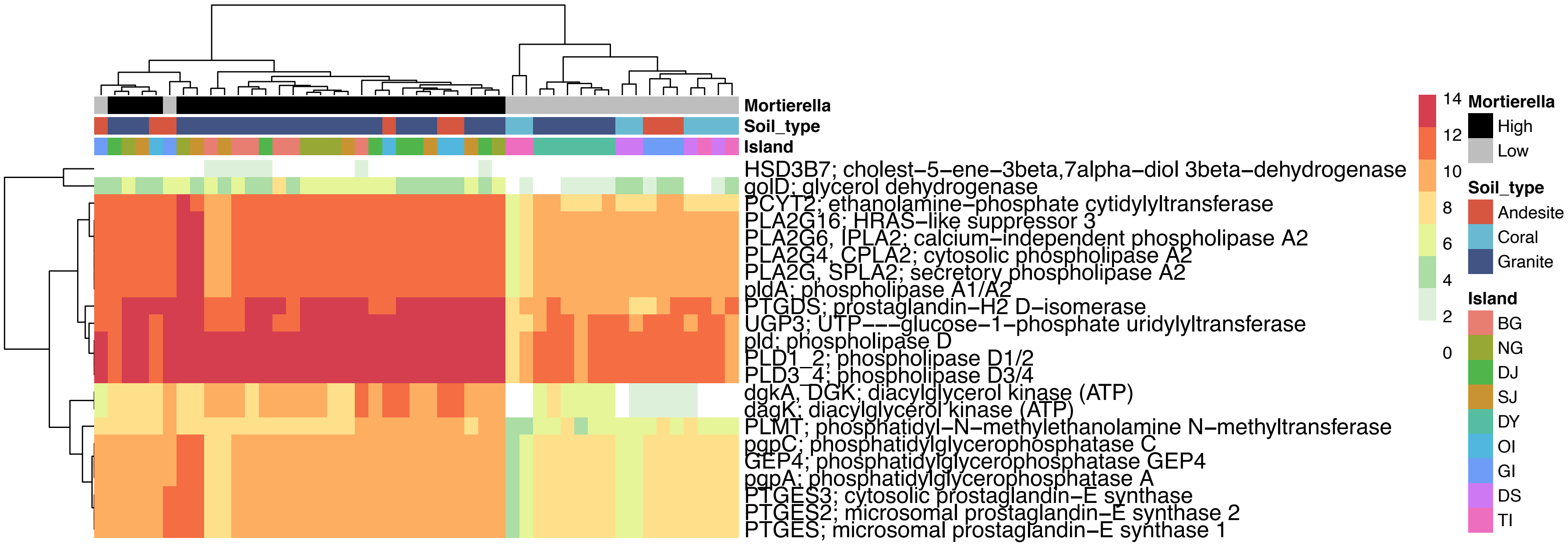

### Supplementary Tables

**Supplementary Table 1.** Sequencing and genome-assembly statistics of *Mortierella* isolates and the representative species from JGI Mycocosm database used in the improvement of PICRUSt2 fungal database

**Supplementary Table 1 is a xlsx file**

**Supplementary Table 2.** Alpha diversity indices of the offshore islands' fungal communities. [a,b,c] Denotes significant differences in the median alpha diversity indices between the islands.

| Island | Median Chao1 ± SD | Median ENS ± SD |
|---|---|---|
| **MT-BG** | 2035.0 ± 570.4[c] | 66.8 ± 94.3[ab] |
| **MT-NG** | 1719.5 ± 333.2[c] | 53.1 ± 35.8[a] |
| **MT-DJ** | 1809.3 ± 402.1[c] | 56.9 ± 25.4[a] |
| **MT-SJ** | 1910.2 ± 266.9[c] | 43.6 ± 20.7[a] |
| **MT-DY** | 864.0 ± 56.2[ab] | 61.4 ± 13.1[a] |
| **OI** | 1323.2 ± 577.4[bc] | 47.3 ± 53.7[ab] |
| **GI** | 1850.7 ±629.1[c] | 142.0 ±100.1[b] |
| **DS** | 976.4 ±31.7[ab] | 88.0 ± 24.7[ab] |
| **TP** | 373.6 ±122.4[a] | 35.5 ± 13.5[a] |

**Supplementary Table 3**. Edaphic and climatic metadata of the nine offshore islands.
ANOVA significant differences between islands are marked using asterisks (*).
[a]Phosphate concentration of coral islands exceeded the assay's detection limit. [b]No weather station data were available.

| | **Beigan** | **Dongju** | **Nangan** | **Hsiju** | **Dongyin** | **Orchid Island** | **Green Island** | **Dongsha islet** | **Taiping islet** |
|---|---|---|---|---|---|---|---|---|---|
| **Abbreviation** | MT-BG | MT-DJ | MT-NG | MT-SJ | MT-DY | OI | GI | DS | TP |
| **Longitude** | 119.989 | 119.970 | 119.933 | 119.939 | 120.496 | 121.548 | 121.490 | 116.732 | 114.365 |
| **Latitude** | 26.223 | 25.952 | 26.150 | 25.974 | 26.371 | 22.045 | 22.664 | 20.701 | 10.377 |
| **Collection date** | 2016/10 | 2016/10 | 2016/10 | 2016/10 | 2017/04 | 2016/11 | 2018/02 | 2017/07 | 2018/07 |
| **Island type** | Mainland | Mainland | Mainland | Mainland | Mainland | Volcanic | Volcanic | Coral | Coral |
| **Rock type** | Granite | Granite | Granite | Granite | Granite | Andesite | Andesite | Coral | Coral |
| **Soil type** | Haplustult | Haplustults | Haplustults | Haplustult | Haplustult | Paleudult | Paleudult | Entisol | Entisol |
| **Island size (km$^2$)** | 9.3 | 2.64 | 10.64 | 2.37 | 4.4 | 45 | 15.09 | 1.8 | 0.51 |
| ***pH** | 4.39 | 4.76 | 4.91 | 4.66 | 4.74 | 6.10 | 7.60 | 8.07 | 7.58 |
| ***MBC** | 551.76 | 432.53 | 470.70 | 492.72 | 550.44 | 1241.57 | 1115.79 | 1156.05 | 1155.15 |
| ***MBN** | 78.92 | 67.41 | 77.93 | 78.03 | 92.51 | 234.21 | 143.24 | 117.48 | 229.51 |
| ***MBP** | 22.27 | 23.41 | 12.14 | 17.51 | 13.32 | 12.11 | 8.86 | [a]NA | [a]NA |
| ***Cellulase** | 2294.94 | 1013.30 | 832.77 | 701.94 | 692.60 | 960.09 | 834.44 | 450.25 | 525.04 |
| ***Xylanase** | 5112.48 | 2931.88 | 2842.81 | 2294.22 | 2922.98 | 2529.75 | 2859.74 | 2195.52 | 2277.54 |
| **Phosphomonoesterase** | 1542.58 | 867.94 | 1509.98 | 849.23 | 551.45 | 2762.18 | 437.28 | 217.73 | 304.71 |
| **Urease** | 1.65 | 1.27 | 1.01 | 0.67 | 2.08 | 10.48 | 2.75 | 3.63 | 5.16 |
| **Proteinase** | 189.12 | 152.79 | 91.86 | 121.39 | 134.45 | 180.81 | 487.52 | 301.83 | 590.80 |
| ***B.glucosaminidase** | 339.20 | 215.72 | 243.55 | 226.68 | 81.53 | 340.89 | 49.43 | 35.37 | 134.56 |
| ***PLFA-total** | 37.20 | 22.70 | 23.00 | 30.20 | 48.70 | 50.60 | 75.12 | 38.85 | NA |
| **PLFA-B** | 15.90 | 9.92 | 9.43 | 13.70 | 21.60 | 23.70 | 34.96 | 17.90 | NA |
| ***PLFA-F** | 0.95 | 0.45 | 0.97 | 0.68 | 1.66 | 0.75 | 2.07 | 0.79 | NA |
| ***PLFA-AMF** | 1.14 | 0.62 | 0.72 | 1.01 | 1.44 | 1.08 | 3.07 | 1.24 | NA |
| **MMT** | 23.80 | 24.20 | 23.80 | 24.20 | [b]NA | 22.20 | 19.10 | [b]NA | [b]NA |
| **MMP** | 40.20 | [b]NA | 40.20 | [b]NA | [b]NA | 385.30 | 22.00 | [b]NA | [b]NA |
| **Vegetation** | Broadleaf forest | Broadleaf forest | Broadleaf forest | Broadleaf forest | *Pinus thunbergii* / *Pittosporum tobira* / *Acacia confusa* | Broadleaf forest | *Pandanus tectorius* | *Pandanus tectorius* | *Terminalia catappa* |

**Supplementary Table 4**. The 56 cosmopolitan zOTUs found across the offshore islands of Taiwan

| ZOTUs | Phylum | Class | Order | Family | Genus | Species |
|---|---|---|---|---|---|---|
| Zotu1 | Mortierellomycota | Mortierellomycetes | Mortierellales | Mortierellaceae | Mortierella | unidentified |
| Zotu2 | Ascomycota | Sordariomycetes | Hypocreales | Nectriaceae | Fusarium | falciforme |
| Zotu3 | Basidiomycota | Agaricomycetes | Hymenochaetales | Hymenochaetaceae | Pyrrhoderma | hainanense |
| Zotu7 | Mortierellomycota | Mortierellomycetes | Mortierellales | Mortierellaceae | Mortierella | unidentified |
| Zotu8 | Basidiomycota | Agaricomycetes | Agaricales | Mycenaceae | Mycena | fulgoris |
| Zotu15 | Mortierellomycota | Mortierellomycetes | Mortierellales | Mortierellaceae | Mortierella | unidentified |
| Zotu17 | Basidiomycota | Tremellomycetes | Tremellales | Trimorphomycetaceae | Saitozyma | podzolica |
| Zotu21 | Ascomycota | Sordariomycetes | Hypocreales | Bionectriaceae | Clonostachys | unidentified |
| Zotu27 | Ascomycota | Sordariomycetes | Hypocreales | Hypocreaceae | Trichoderma | unidentified |
| Zotu29 | Ascomycota | Sordariomycetes | Hypocreales | Nectriaceae | Pleiocarpon | algeriense |
| Zotu42 | Ascomycota | Sordariomycetes | Hypocreales | Nectriaceae | Fusarium | foetens |
| Zotu44 | Ascomycota | Sordariomycetes | Hypocreales | Nectriaceae | Fusarium | obliquiseptatum |
| Zotu45 | Ascomycota | Sordariomycetes | Hypocreales | Nectriaceae | Neocosmospora | perseae |
| Zotu46 | Ascomycota | Sordariomycetes | Hypocreales | Bionectriaceae | Clonostachys | solani |
| Zotu48 | Ascomycota | Sordariomycetes | Hypocreales | Hypocreaceae | Trichoderma | yunnanense |
| Zotu67 | Ascomycota | Sordariomycetes | Hypocreales | Clavicipitaceae | Metarhizium | unidentified |
| Zotu69 | Ascomycota | Sordariomycetes | Hypocreales | Clavicipitaceae | Metarhizium | cylindrosporum |
| Zotu70 | Ascomycota | Sordariomycetes | Hypocreales | Nectriaceae | Fusarium | unidentified |
| Zotu73 | Ascomycota | Sordariomycetes | Sordariales | Chaetomiaceae | Humicola | homopilata |
| Zotu75 | Ascomycota | Sordariomycetes | Hypocreales | Hypocreaceae | Trichoderma | crassum |
| Zotu87 | Mortierellomycota | Mortierellomycetes | Mortierellales | Mortierellaceae | Mortierella | SH1557014.08FU |
| Zotu96 | Ascomycota | Sordariomycetes | Hypocreales | Clavicipitaceae | Metarhizium | unidentified |
| Zotu115 | Ascomycota | Dothideomycetes | Botryosphaeriales | Botryosphaeriaceae | Lasiodiplodia | unidentified |
| Zotu118 | Ascomycota | Sordariomycetes | Hypocreales | Nectriaceae | Fusarium | unidentified |
| Zotu119 | Ascomycota | Sordariomycetes | Hypocreales | Clavicipitaceae | Metacordyceps | unidentified |
| Zotu121 | Ascomycota | Sordariomycetes | Sordariales | Chaetomiaceae | Humicola | olivacea |
| Zotu163 | Ascomycota | Sordariomycetes | Sordariales | sedis | Staphylotrichum | SH1615643.08FU |
| Zotu172 | Ascomycota | Sordariomycetes | Hypocreales | Hypocreaceae | Trichoderma | rugulosum |
| Zotu177 | Ascomycota | Eurotiomycetes | Eurotiales | Aspergillaceae | Aspergillus | unidentified |
| Zotu182 | Ascomycota | Sordariomycetes | Hypocreales | Clavicipitaceae | Metacordyceps | unidentified |
| Zotu189 | Ascomycota | Sordariomycetes | Xylariales | Sporocadaceae | Pestalotiopsis | unidentified |
| Zotu201 | Ascomycota | Sordariomycetes | Sordariales | Chaetomiaceae | Humicola | homopilata |
| Zotu205 | Ascomycota | Sordariomycetes | Hypocreales | Ophiocordycipitaceae | Purpureocillium | lavendulum |
| Zotu206 | Ascomycota | Sordariomycetes | Hypocreales | Nectriaceae | Fusarium | unidentified |
| Zotu226 | Ascomycota | Sordariomycetes | Hypocreales | Bionectriaceae | Clonostachys | unidentified |
| Zotu255 | Ascomycota | Sordariomycetes | Trichosphaeriales | Trichosphaeriaceae | Nigrospora | SH1549625.08FU |
| Zotu258 | Ascomycota | Dothideomycetes | Cladosporiales | Cladosporiaceae | Cladosporium | oxysporum |
| Zotu299 | Basidiomycota | Agaricomycetes | Agaricales | Clavariaceae | Ramariopsis | flavescens |
| Zotu304 | Ascomycota | Sordariomycetes | Sordariales | sedis | Staphylotrichum | unidentified |
| Zotu333 | Ascomycota | Eurotiomycetes | Eurotiales | Aspergillaceae | Aspergillus | unidentified |
| Zotu456 | Ascomycota | Dothideomycetes | Capnodiales | Mycosphaerellaceae | Phaeophleospora | SH1606643.08FU |
| Zotu602 | Ascomycota | Dothideomycetes | Capnodiales | Capnodiaceae | Leptoxyphium | SH1714997.08FU |
| Zotu607 | Ascomycota | Eurotiomycetes | Eurotiales | Trichocomaceae | Talaromyces | unidentified |
| Zotu630 | Basidiomycota | Agaricomycetes | Amylocorticiales | Amylocorticiaceae | Podoserpula | ailaoshanensis |
| Zotu654 | Ascomycota | Dothideomycetes | Pleosporales | Didymellaceae | Allophoma | cylindrispora |
| Zotu663 | Ascomycota | Sordariomycetes | Glomerellales | Plectosphaerellaceae | Chordomyces | SH1644390.08FU |
| Zotu745 | Ascomycota | Sordariomycetes | Hypocreales | Bionectriaceae | Nectriopsis | lindauiana |
| Zotu775 | Ascomycota | Dothideomycetes | Pleosporales | Didymellaceae | Stagonosporopsis | lupini |
| Zotu796 | Ascomycota | Sordariomycetes | Glomerellales | Plectosphaerellaceae | Chordomyces | antarcticum |
| Zotu908 | Ascomycota | Dothideomycetes | Capnodiales | Cladosporiaceae | Cladosporium | unidentified |
| Zotu1150 | Ascomycota | Dothideomycetes | Pleosporales | Thyridariaceae | Roussoella | SH1637047.08FU |
| Zotu2857 | Ascomycota | Leotiomycetes | Helotiales | Hyaloscyphaceae | Hyphodiscus | brevicollaris |
| Zotu2912 | Ascomycota | Dothideomycetes | Pleosporales | Didymellaceae | Stagonosporopsis | lupini |
| Zotu4316 | Ascomycota | Sordariomycetes | Hypocreales | Nectriaceae | Volutella | unidentified |
| Zotu5550 | Ascomycota | Sordariomycetes | Hypocreales | Bionectriaceae | Clonostachys | SH1155568.08FU |
| Zotu7679 | Basidiomycota | Malasseziomycetes | Malasseziales | Malasseziaceae | Malassezia | unidentified |

**Supplementary Table 5.** Pairwise Tukey post hoc significant test between the degreeness of different fungal phyla in each island. Significant difference with p-values less than 0.05 are coloured in red.

| Island : MT-BG | Mortierellomycota | Mucoromycota | Ascomycota | Basidiomycota |
|---|---|---|---|---|
| Mucoromycota | 0.991 | | | |
| Ascomycota | 0.735 | 0.222 | | |
| Basidiomycota | 0.964 | 0.636 | 0.853 | |
| Chytridiomycota | 1.000 | 0.993 | 0.807 | 0.975 |

| Island : OI | Mortierellomycota | Mucoromycota | Ascomycota | Basidiomycota |
|---|---|---|---|---|
| Mucoromycota | 0.328 | | | |
| Ascomycota | 0.293 | 0.965 | | |
| Basidiomycota | 0.132 | 1.000 | 0.998 | |
| Chytridiomycota | 0.101 | 0.900 | 0.518 | 0.775 |

| Island : MT-NG | Mortierellomycota | Mucoromycota | Ascomycota | Basidiomycota |
|---|---|---|---|---|
| Mucoromycota | 0.011 | | | |
| Ascomycota | 0.037 | 0.699 | | |
| Basidiomycota | 0.001 | 0.999 | 0.164 | |
| Chytridiomycota | 0.000 | 0.426 | 0.041 | 0.433 |

| Island : GI | Mortierellomycota | Mucoromycota | Ascomycota | Basidiomycota |
|---|---|---|---|---|
| Mucoromycota | 0.587 | | | |
| Ascomycota | 0.140 | 0.996 | | |
| Basidiomycota | 0.204 | 0.999 | 0.999 | |
| Chytridiomycota | 0.378 | 1.000 | 0.999 | 1.000 |

| Island : MT-DJ | Mortierellomycota | Mucoromycota | Ascomycota | Basidiomycota |
|---|---|---|---|---|
| Mucoromycota | 0.712 | | | |
| Ascomycota | 0.788 | 0.982 | | |
| Basidiomycota | 0.862 | 0.976 | 1.000 | |
| Chytridiomycota | 0.654 | 1.000 | 0.959 | 0.951 |

| Island : DS | Mortierellomycota | Mucoromycota | Ascomycota | Basidiomycota |
|---|---|---|---|---|
| Mucoromycota | 0.040 | | | |
| Ascomycota | 0.712 | 0.022 | | |
| Basidiomycota | 0.634 | 0.088 | 0.990 | |
| Chytridiomycota | 0.348 | 0.768 | 0.672 | 0.862 |

| Island : MT-SJ | Mortierellomycota | Mucoromycota | Ascomycota | Basidiomycota |
|---|---|---|---|---|
| Mucoromycota | 0.271 | | | |
| Ascomycota | 0.113 | 0.999 | | |
| Basidiomycota | 0.154 | 0.999 | 1.000 | |
| Chytridiomycota | 0.964 | 0.850 | 0.842 | 0.861 |

| Island : TP | Mortierellomycota | Mucoromycota | Ascomycota | Basidiomycota |
|---|---|---|---|---|
| Mucoromycota | 0.950 | | | |
| Ascomycota | 0.985 | 0.976 | | |
| Basidiomycota | 0.945 | 1.000 | 0.958 | |
| Chytridiomycota | 0.998 | 0.976 | 0.998 | 0.965 |

| Island : MT-DY | Mortierellomycota | Mucoromycota | Ascomycota | Basidiomycota |
|---|---|---|---|---|
| Mucoromycota | 0.998 | | | |
| Ascomycota | 0.984 | 1.000 | | |
| Basidiomycota | 0.999 | 0.955 | 0.438 | |
| Chytridiomycota | 0.999 | 1.000 | 0.998 | 0.967 |

**Supplementary Table 6. Pairwise Tukey post hoc significant test between the mean edge weight of different fungal phyla in each island. Significant difference with p-values less than 0.05 are coloured in red.**

| Island : MT-BG | Mortierellomycota | Mucoromycota | Ascomycota | Basidiomycota |
|---|---|---|---|---|
| Mucoromycota | 0.000 | | | |
| Ascomycota | 0.000 | 0.340 | | |
| Basidiomycota | 0.000 | 0.645 | 0.968 | |
| Chytridiomycota | 0.000 | 0.351 | 0.917 | 0.818 |

| Island : OI | Mortierellomycota | Mucoromycota | Ascomycota | Basidiomycota |
|---|---|---|---|---|
| Mucoromycota | 0.006 | | | |
| Ascomycota | 0.000 | 1.000 | | |
| Basidiomycota | 0.000 | 1.000 | 1.000 | |
| Chytridiomycota | 0.003 | 0.820 | 0.794 | 0.814 |

| Island : MT-NG | Mortierellomycota | Mucoromycota | Ascomycota | Basidiomycota |
|---|---|---|---|---|
| Mucoromycota | 0.040 | | | |
| Ascomycota | 0.000 | 0.419 | | |
| Basidiomycota | 0.000 | 0.439 | 1.000 | |
| Chytridiomycota | 0.064 | 0.987 | 0.984 | 0.979 |

| Island : GI | Mortierellomycota | Mucoromycota | Ascomycota | Basidiomycota |
|---|---|---|---|---|
| Mucoromycota | 0.637 | | | |
| Ascomycota | 0.019 | 0.667 | | |
| Basidiomycota | 0.005 | 0.354 | 0.674 | |
| Chytridiomycota | 0.006 | 0.271 | 0.612 | 0.973 |

| Island : MT-DJ | Mortierellomycota | Mucoromycota | Ascomycota | Basidiomycota |
|---|---|---|---|---|
| Mucoromycota | 0.034 | | | |
| Ascomycota | 0.001 | 1.000 | | |
| Basidiomycota | 0.001 | 0.989 | 0.981 | |
| Chytridiomycota | 0.077 | 1.000 | 0.995 | 0.966 |

| Island : DS | Mortierellomycota | Mucoromycota | Ascomycota | Basidiomycota |
|---|---|---|---|---|
| Mucoromycota | 0.103 | | | |
| Ascomycota | 0.376 | 0.493 | | |
| Basidiomycota | 0.181 | 0.928 | 0.732 | |
| Chytridiomycota | 0.137 | 1.000 | 0.627 | 0.965 |

| Island : MT-SJ | Mortierellomycota | Mucoromycota | Ascomycota | Basidiomycota |
|---|---|---|---|---|
| Mucoromycota | 0.022 | | | |
| Ascomycota | 0.000 | 0.030 | | |
| Basidiomycota | 0.000 | 0.151 | 0.931 | |
| Chytridiomycota | 0.000 | 0.126 | 0.973 | 0.877 |

| Island : TP | Mortierellomycota | Mucoromycota | Ascomycota | Basidiomycota |
|---|---|---|---|---|
| Mucoromycota | 0.994 | | | |
| Ascomycota | 1.000 | 0.882 | | |
| Basidiomycota | 1.000 | 0.988 | 0.952 | |
| Chytridiomycota | 0.993 | 1.000 | 0.851 | 0.985 |

| Island : MT-DY | Mortierellomycota | Mucoromycota | Ascomycota | Basidiomycota |
|---|---|---|---|---|
| Mucoromycota | 0.916 | | | |
| Ascomycota | 0.999 | 0.521 | | |
| Basidiomycota | 0.996 | 0.478 | 0.999 | |
| Chytridiomycota | 0.999 | 0.959 | 0.950 | 0.920 |

**Supplementary Table 7.** The 45 hub fungal zOTUs determined from different offshore island of Taiwan. Cosmopolitan taxa are indicated by the 'Ubiquitous' column.

| zOTU | Phylum | Class | Order | Family | Genus | Species | Node Type | Ubiquitous |
|---|---|---|---|---|---|---|---|---|
| Zotu1 | Mortierellomycota | Mortierellomycetes | Mortierellales | Mortierellaceae | Mortierella | unidentified | Module hubs | Yes |
| Zotu7 | Mortierellomycota | Mortierellomycetes | Mortierellales | Mortierellaceae | Mortierella | unidentified | Module hubs | Yes |
| Zotu21 | Ascomycota | Sordariomycetes | Hypocreales | Bionectriaceae | Clonostachys | unidentified | Module hubs | Yes |
| Zotu75 | Ascomycota | Sordariomycetes | Hypocreales | Hypocreaceae | Trichoderma | crassum | Network hubs | Yes |
| Zotu226 | Ascomycota | Sordariomycetes | Hypocreales | Bionectriaceae | Clonostachys | unidentified | Module hubs | Yes |
| Zotu304 | Ascomycota | Sordariomycetes | Sordariales | sedis | Staphylotrichum | unidentified | Module hubs | Yes |
| Zotu333 | Ascomycota | Eurotiomycetes | Eurotiales | Aspergillaceae | Aspergillus | unidentified | Module hubs | Yes |
| Zotu5 | Mortierellomycota | Mortierellomycetes | Mortierellales | Mortierellaceae | Mortierella | unidentified | Module hubs | No |
| Zotu13 | Mortierellomycota | Mortierellomycetes | Mortierellales | Mortierellaceae | Mortierella | unidentified | Network hubs | No |
| Zotu16 | Basidiomycota | Agaricomycetes | Agaricales | Hygrophoraceae | Hygrocybe | roseopallida | Module hubs | No |
| Zotu43 | Ascomycota | Saccharomycetes | Saccharomycetales | Trichomonascaceae | Blastobotrys | serpentis | Module hubs | No |
| Zotu52 | Basidiomycota | Tremellomycetes | Tremellales | Trimorphomycetaceae | Saitozyma | podzolica | Network hubs | No |
| Zotu57 | Basidiomycota | Agaricomycetes | Agaricales | Entolomataceae | Clitopilus | unidentified | Network hubs | No |
| Zotu60 | Ascomycota | Sordariomycetes | Hypocreales | Nectriaceae | Gliocephalotrichum | humicola | Network hubs | No |
| Zotu63 | Ascomycota | Leotiomycetes | Helotiales | Helotiaceae | Tetracladium | unidentified | Module hubs | No |
| Zotu71 | Basidiomycota | Agaricomycetes | Agaricales | Entolomataceae | Entoloma | nigrovelutinum | Module hubs | No |
| Zotu82 | Mortierellomycota | Mortierellomycetes | Mortierellales | Mortierellaceae | Mortierella | unidentified | Module hubs | No |
| Zotu91 | Ascomycota | Dothideomycetes | Pleosporales | Testudinaceae | Neotestudina | rosatii | Network hubs | No |
| Zotu109 | Chytridiomycota | Chytridiomycetes | Rhizophydiales | Protrudomycetaceae | Protrudomyces | lateralis | Module hubs | No |
| Zotu131 | Basidiomycota | Agaricomycetes | Trechisporales | Hydnodontaceae | Trechispora | unidentified | Network hubs | No |
| Zotu218 | Basidiomycota | Agaricomycetes | Agaricales | Agaricaceae | Leucoagaricus | variicolor | Module hubs | No |
| Zotu245 | Ascomycota | Eurotiomycetes | Onygenales | Ajellomycetaceae | Blastomyces | helicus | Module hubs | No |
| Zotu276 | Ascomycota | Dothideomycetes | Pleosporales | unidentified | Pseudoasteromassaria | fagi | Network hubs | No |
| Zotu288 | Chytridiomycota | Chytridiomycetes | Lobulomycetales | Lobulomycetaceae | Clydaea | vesicula | Network hubs | No |
| Zotu368 | Ascomycota | Dothideomycetes | unidentified | unidentified | Soloacrosporiella | acaciae | Network hubs | No |
| Zotu369 | Mortierellomycota | Mortierellomycetes | Mortierellales | Mortierellaceae | Mortierella | unidentified | Module hubs | No |
| Zotu386 | Ascomycota | Dothideomycetes | Pleosporales | unidentified | unidentified | sp. | Network hubs | No |
| Zotu430 | Ascomycota | Geoglossomycetes | Geoglossales | Geoglossaceae | Geoglossum | chamaecyparinum | Network hubs | No |
| Zotu454 | Ascomycota | Sordariomycetes | Hypocreales | Hypocreaceae | Trichoderma | unidentified | Network hubs | No |
| Zotu482 | Basidiomycota | Agaricomycetes | Trechisporales | Hydnodontaceae | Trechispora | unidentified | Module hubs | No |
| Zotu545 | Basidiomycota | Agaricomycetes | Russulales | Russulaceae | Russula | unidentified | Module hubs | No |
| Zotu664 | Ascomycota | Sordariomycetes | Sordariales | Chaetomiaceae | Podospora | dimorpha | Network hubs | No |
| Zotu686 | Ascomycota | Sordariomycetes | Sordariales | Lasiosphaeriaceae | Apodus | deciduus | Network hubs | No |
| Zotu902 | Chytridiomycota | Chytridiomycetes | Rhizophydiales | unidentified | Coralloidiomyces | digitatus | Network hubs | No |
| Zotu940 | Ascomycota | Dothideomycetes | unidentified | unidentified | Marquesius | aquaticus | Network hubs | No |
| Zotu1127 | Basidiomycota | Agaricomycetes | Agaricales | Agaricaceae | Coprinus | trigonosporus | Network hubs | No |
| Zotu1237 | Ascomycota | Eurotiomycetes | Sclerococcales | Dactylosporaceae | Sclerococcum | vrijmoediae | Network hubs | No |
| Zotu1583 | Basidiomycota | Microbotryomycetes | unidentified | Chrysozymaceae | Sampaiozyma | vanillica | Network hubs | No |
| Zotu1725 | Basidiomycota | Agaricomycetes | Agaricales | Agaricaceae | Macrolepiota | orientiexcoriata | Network hubs | No |
| Zotu2123 | Ascomycota | unidentified | unidentified | unidentified | Pseudopenidiella | piceae | Network hubs | No |
| Zotu2219 | Ascomycota | Orbiliomycetes | Orbiliales | Orbiliaceae | Drechslerella | dactyloides | Network hubs | No |
| Zotu2265 | Mucoromycota | Mortierellomycetes | Mortierellales | Mortierellaceae | Mortierella | globulifera | Network hubs | No |
| Zotu2518 | Ascomycota | Geoglossomycetes | Geoglossales | Geoglossaceae | Geoglossum | variabilisporum | Network hubs | No |
| Zotu4549 | Basidiomycota | Agaricomycetes | Sebacinales | Sebacinaceae | Sebacina | cystidiata | Network hubs | No |

**Supplementary Table 8**. Differentially predicted pathway and KO using original and improved PICRUSt2 fungal database

| | **Differential predicted terms with log fold change greater than 2** | | | |
|---|---|---|---|---|
| **PICRUSt2 database** | **Pathways Up** | **Pathways Down** | **KO Up** | **KO Down** |
| **Original** | 1 | 3 | 25 | 19 |
| **Improved** | 3 | 1 | 243 | 133 |
