## Supplementary figures and images for "Genomes of keystone *Mortierella* species lead to better *in silico* prediction of soil mycobiome functions from Taiwan’s offshore islands"

### Supplementary Fig 5

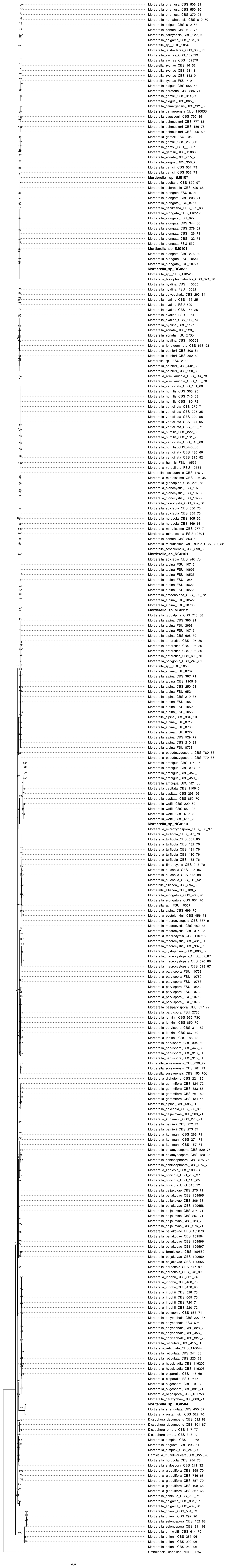
